# Leveraging Multimodal Large Language Models to Extract Mechanistic Insights from Biomedical Visuals: A Case Study on COVID-19 and Neurodegenerative Diseases

**DOI:** 10.1101/2025.10.01.679928

**Authors:** Elizaveta Popova, Marc Jacobs, Martin Hofmann-Apitius, Negin Sadat Babaiha

## Abstract

**Background:** The COVID-19 pandemic has intensified concerns about its long-term neurological impact, with growing evidence linking SARS-CoV-2 infection to neurodegenerative diseases (NDDs) such as Alzheimer’s (AD) and Parkinson’s (PD). Patients with these conditions not only face higher risk of severe COVID-19 outcomes but may also undergo accelerated cognitive and motor decline following infection. Proposed mechanisms—ranging from neuroinflammation and blood–brain barrier disruption to abnormal protein aggregation—closely mirror core features of neurodegenerative pathology. Yet, current knowledge is fragmented across text, figures, and pathway diagrams, hindering integration into computational models capable of uncovering systemic patterns.

**Results:** To address this gap, we applied GPT-4 Omni (GPT-4o), a multimodal large language model, to extract mechanistic insights from biomedical figures. Over 10,000 images were retrieved through targeted searches on COVID-19 and neurodegeneration; after automated and manual filtering, a curated subset was analyzed. GPT-4o extracted biological relationships as semantic triples, which were grouped into six mechanistic categories—including microglial activation and barrier disruption—using ontology-guided similarity and assembled into a Neo4j knowledge graph.

Accuracy was evaluated against a gold-standard dataset of expert-annotated images using BioBERT-based semantic matching. This evaluation also enabled prompt tuning, threshold optimization, and hyperparameter assessment. Results demonstrate that GPT-4o successfully recovers both established and novel mechanisms, yielding interpretable outputs that illuminate complex biological links between SARS-CoV-2 and neurodegeneration.

**Conclusions:** This study showcases the potential of multimodal LLMs to mine biomedical visual data at scale. By complementing text mining and integrating figure-derived knowledge, our framework advances understanding of COVID-19–related neurodegeneration and supports future translational research.

## 1. Introduction

The outbreak of the COVID-19 pandemic, caused by the coronavirus SARS-CoV-2, has led to a global health crisis of unprecedented scale and complexity since its identification in late 2019 (Fu et al., 2022; C. Huang et al., 2023; C. Li et al., 2022; J. Li et al., 2021; L. Mao et al., 2020). While much of the early focus was directed at the acute respiratory effects of the virus, it quickly became evident that SARS-CoV-2 infection can result in systemic complications, persisting well beyond the resolution of primary symptoms. Among patients who have recovered from the initial infection, a wide range of long-term effects — collectively referred to as long COVID — have been reported. These include chronic fatigue, reduced pulmonary function, renal dysfunction, cardiovascular irregularities, and neurological symptoms such as brain fog, memory impairment, and cognitive decline (Fu et al., 2022; Heneka et al., 2020; C. Huang et al., 2023; C. Li et al., 2022; Mahalakshmi et al., 2021; L. Mao et al., 2020).

Of particular concern is the mounting evidence that COVID-19 may influence or exacerbate NDDs such as AD and PD (De Felice et al., 2020; Heneka et al., 2020; Mahalakshmi et al., 2021; Wu et al., 2020). The virus’s ability to trigger neuroinflammatory processes and disrupt the blood-brain barrier, alongside documented cases of microglial activation and tau phosphorylation, has fueled speculation that SARS-CoV-2 may accelerate preclinical neurodegeneration or initiate novel pathological cascades. Although these findings are suggestive, our understanding of the causal and mechanistic links between COVID-19 and neurodegeneration remains limited.

Observational and epidemiological studies are constrained by confounding factors — age, infection severity, hospitalization status, and comorbidities such as hypertension or cardiovascular disease — making it difficult to isolate the specific contribution of SARS-CoV-2 to long-term neurological outcomes (Mizrahi et al., 2023; Xu et al., 2022). Furthermore, these studies are often retrospective and lack molecular resolution, limiting their utility for uncovering underlying biological mechanisms. As a result, there is a growing need for integrative approaches that can connect clinical observations to mechanistic insights, potentially guiding the development of targeted interventions.

One promising but underutilized avenue lies in the systematic analysis of biomedical figures, particularly graphical abstracts, signaling pathway diagrams, and schematic illustrations, which frequently distill complex biological relationships into intuitive visual representations. These figures are not mere decorative elements; rather, they function as condensed knowledge units, often revealing spatial, temporal, or regulatory dynamics that are difficult to capture through narrative text alone. For example, pathway diagrams showing the cascade from ACE2 receptor binding to neuroinflammatory cytokine release illustrate multi-step biological processes that would otherwise require dense, linear descriptions.

Despite their value, such figures are rarely included in literature mining workflows, which remain largely text-centric (Babaiha et al., 2023, 2024). Traditional search platforms like PubMed index primarily textual metadata — titles, abstracts, keywords — thereby omitting papers that contribute valuable visual content. As a result, a substantial portion of scientific knowledge encoded in figures remains inaccessible to automated analysis and downstream applications such as KG construction, hypothesis generation, and mechanistic modeling (Ahmed et al., 2016; Babaiha et al., 2025; Trelles Trabucco et al., 2023).

Recent developments in multimodal artificial intelligence (AI) — particularly unified vision language models (VLMs) — have opened new possibilities for integrating visual content into scientific data mining (Medical Vision Language Pretraining: A Survey, n.d.). VLMs are designed to jointly encode and reason over both visual and textual inputs, enabling them to interpret complex image-based content in a manner that approximates human multimodal understanding. These models are typically trained on large-scale paired datasets of images and captions, allowing them to align visual elements (e.g., molecular pathways, cellular diagrams, microscopy) with linguistic concepts (Liu et al., 2023). For example, BiomedGPT, a domain-adapted VLM, has demonstrated strong performance across a range of biomedical tasks, including radiology report generation, dermatology image captioning, and visual question answering in pathology settings (Zhang et al., 2024). Its architecture integrates medical image understanding with language generation capabilities, enabling it to bridge diagnostic visuals and clinical narratives effectively. Similarly, GPT-4V, the vision-augmented extension of GPT-4, has shown impressive generalization in biomedical image interpretation tasks such as histopathological classification, where it achieves performance comparable to that of domain-specific convolutional neural networks—even in few-shot, in-context learning settings (Ono et al., 2024).

The clinical utility of VLMs has also been increasingly validated through real-world medical evaluation. In a notable study, Han et al. (Han et al., 2023) assessed GPT-4V on complex clinical case questions sourced from high-impact medical journals such as the Journal of the American Medical Association and the New England Journal of Medicine. The model successfully synthesized multimodal input — including patient histories and diagnostic images — and outperformed both human respondents and text-only LLMs, highlighting its potential as a decision-support tool in clinical reasoning. Similarly, Dong et al. (Dong et al., 2024) applied GPT-4V to the task of generating pathology reports from histopathological images, showing that the model could produce biologically detailed and diagnostically meaningful captions. These outputs were not only coherent and clinically relevant but also enhanced interpretability, offering promise for automating components of clinical documentation and augmenting diagnostic workflows.

These advances in multimodal modeling are unfolding alongside a parallel evolution in KG construction, offering synergistic opportunities for structured biomedical reasoning. KGs encode scientific knowledge as directed graphs, where nodes represent entities such as proteins, diseases, or biological processes, and edges capture their relationships — such as “inhibits,” “activates,” or “expressed in”. When effectively built and curated, KGs serve as dynamic, extensible frameworks for integrating heterogeneous data types and uncovering non-obvious associations across biological scales (Ehrlinger & Wöß, 2016; Hogan et al., 2021).

The utility of KGs in biomedical research is well-established. For example, the SPOKE graph integrates millions of biomedical concepts — including genes, drugs, symptoms, and diseases — into a unified, queryable structure that supports translational insights and systems-level hypothesis generation (Soman et al., 2024). In parallel, Machine Learning (ML)-powered tools like SciSpacy and PubTator have shown that structured triples can be reliably extracted from biomedical literature to populate KGs, enabling downstream applications in semantic search, knowledge discovery, and automated reasoning (Kilicoglu et al., 2011; Neumann et al., 2019; Wei et al., 2024).

These developments underscore a critical shift: from static databases toward computational knowledge infrastructures that can continuously evolve through the integration of both textual and visual data sources — a direction further accelerated by the emergence of VLMs. However, few existing efforts have yet incorporated visual data into this process. This gap is especially glaring given that figures often convey regulatory mechanisms, cellular localization, or cross-talk between pathways — information that is rarely fully replicated in text (Ahmed et al., 2016; Hullman & Bach, 2018). Integrating visual data into information extraction and KG construction holds significant potential for uncovering previously overlooked or underreported biological relationships. Figures often distill complex experimental findings into clear mechanistic representations that are not fully captured in the surrounding text. For example, a graphical abstract may visually depict how SARS-CoV-2 infection triggers NLRP3 inflammasome activation, ultimately leading to neuroinflammation — a mechanistic pathway that may be described ambiguously or omitted entirely from the article’s narrative (Babaiha et al., 2025; Dutta et al., 2022; Guarnieri et al., 2023; Heneka et al., 2020). By extracting such relationships directly from visual content, computational systems can access a richer layer of biological insight, enabling more comprehensive and mechanistically grounded KGs.

To address this limitation, we introduce a novel workflow for extracting semantic triples from biomedical figures — particularly graphical abstracts — using GPT-4o, a state-of-the-art multimodal LLM developed by OpenAI (Han et al., 2023; Liu et al., 2023, p. 4; OpenAI API documentation, 2024). This model is capable of interpreting complex biomedical images and converting them into structured biological statements, which we then use to construct a multimodal KG capturing mechanistic links between COVID-19 and NDD processes.

To construct a representative and diverse corpus of biomedical figures, we leverage Google Image Search as a complementary discovery mechanism. Unlike traditional databases such as PubMed — which are limited to indexing textual content — Google Image Search enables retrieval based on visual context, including diagrammatic style, image captions, and figure structure. This capability supports exploratory, visually driven browsing, allowing researchers to surface semantically relevant figures that might remain undiscovered through conventional keyword-based querying alone (Trelles Trabucco et al., 2023; Yee et al., 2003). By incorporating this modality into our figure selection process, we not only enhance the topical and visual diversity of the corpus but also mirror the real-world strategies used by scientists to navigate, compare, and interpret visual data in biomedical research.

By integrating visual data, multimodal AI, and KG construction, this study introduces a novel, end-to-end framework for mining and structuring mechanistic knowledge from biomedical figures. As a targeted use case, we focus on the emerging intersection and comorbidity between COVID-19 and NDDs — a domain where mechanistic understanding remains evolving. We demonstrate how biomedical figures can surface implicit relationships that are often underreported in textual narratives, including viral modulation of neuronal signaling, glial cell activation, blood-brain barrier disruption, and broader neuroinflammatory cascades (Domingo-Fernández et al., 2021; J. Mao et al., 2019). By extracting and structuring this visual knowledge, the resulting graph captures a richer and more interconnected semantic landscape, enabling downstream applications such as hypothesis generation, multi-scale mechanistic modeling, and exploratory pathways for therapeutic discovery. This case study exemplifies how multimodal approaches can reveal latent biological insight in complex interdisciplinary domains where traditional text mining alone may fall short.

More broadly, this work contributes to the growing recognition that figures are not peripheral illustrations, but primary carriers of scientific knowledge. As multimodal models continue to advance, there is a pressing need for methods that systematically incorporate visual modalities into computational biomedical workflows. Our approach responds to this need by offering a scalable, generalizable strategy for integrating figure-derived knowledge into structured representations — bridging a critical gap between image interpretation and machine-readable scientific reasoning (Ferber et al., 2024; Liu et al., 2023; Medical Vision Language Pretraining: A Survey, n.d.). In doing so, we aim to elevate the role of visual data within the scientific research pipeline, paving the way toward more comprehensive, integrative, and insight-rich biomedical AI systems.

## 2. Materials and methods

### 2.1. Dataset collection

To systematically investigate the comorbidity between COVID-19 and neurodegeneration, we designed a multi-stage pipeline that integrates automated image retrieval, KG construction and analysis. A schematic of the full workflow is illustrated in Fig. 1. The methodology consists of four key components: (1) dataset collection centered on biomedical figures; (2) visual content filtering using an LLM; (3) semantic triple extraction using an LLM; and (4) graph construction, analysis and categorization of extracted biological mechanisms.

**Fig. 1.**
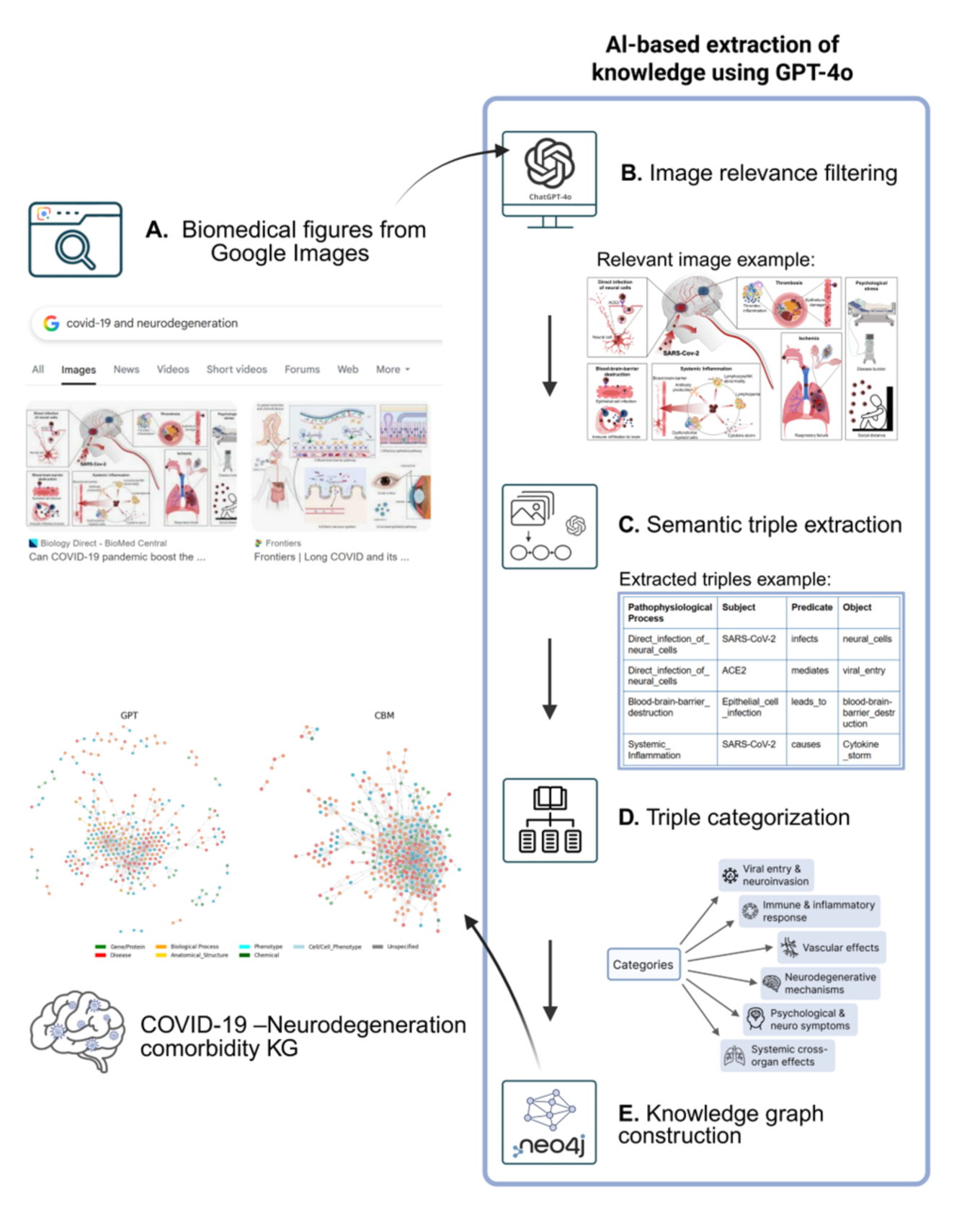
The pipeline for extracting structured mechanistic knowledge from biomedical figures linking COVID-19 and neurodegeneration. (A) Biomedical figures are retrieved from Google Images using the search query “COVID-19 and Neurodegeneration”. (B) Images are filtered for relevance using GPT-4o-based classification and manual review. (C) For each relevant image, semantic triples are extracted, representing various pathophysiological mechanisms. (D) Extracted triples are grouped into six mechanistic categories using ontology-driven semantic similarity. (E) All triples are loaded into a Neo4j database to construct a KG capturing mechanistic links between COVID-19 and neurodegenerative processes and pathways.

To automate the retrieval of image URLs from Google Image Search, we developed a custom web-scraping pipeline using Selenium WebDriver for Chrome browser (Gojare et al., 2015). Selenium enables dynamic interaction with JavaScript-heavy web pages by simulating real-time user behavior, thereby allowing for scalable and reproducible image data collection that closely mimics manual browsing. For this study, we initiated a Google Image search using the query “COVID-19 and neurodegeneration”. The script first extracted the URLs of the top 100 primary images displayed in the search results. To expand the dataset and capture a broader range of visually relevant figures, the script then used Google’s Related Images feature for each of the 100 primary images. From each related image set, 100 visually similar image URLs were extracted, resulting in an initial dataset comprising 10,100 image URLs, which was cleaned and processed during the further steps.

### 2.2. Image relevance assessment

To curate a high-quality corpus of biomedical figures, we applied a multi-stage filtering pipeline combining automated and manual steps. After removing duplicates and inaccessible files, figures were screened for relevance using GPT-4o classification and manual inspection. This process yielded 289 figures depicting mechanistic interactions between COVID-19 and neurodegenerative processes (Table 1). Detailed filtering steps are provided in Supplementary Methods.

**Table 1.** Details of image relevance analysis.

| Category | Number of URLs |
| --- | --- |
| Total URLs | 6319 |
| Validated URLs | 3614 |
| Relevant URLs (GPT), 1st run | 626 |
| Relevant URLs (GPT), 2nd run | 567 |
| Relevant URLs (manual validation) | 289 |

Summary of image relevance filtering pipeline. The table reports the total number of image URLs retrieved, filtered, and validated across multiple stages, including two rounds of automated GPT-4o-based assessment and final manual validation.

### 2.3. Automated triple extraction and evaluation

#### 2.3.1. Pipeline architecture

We developed a custom pipeline utilizing OpenAI’s GPT-4o application programming interface (API) to extract structured biological knowledge from biomedical figures. The pipeline authenticates API access, validates image URL accessibility, and processes each figure to extract biologically meaningful interactions as triples. These triples capture relationships between cells, proteins, small molecules, and biological processes at the intersection of COVID-19 and neurodegenerative mechanisms. Extracted data are parsed into pandas DataFrames and exported in Excel and CSV formats for downstream analysis and KG integration.

#### 2.3.2. Evaluation framework

We collaborated with the Biocuration and knowledge extraction company Causality Biomodels (CBM) (Causality BioModels, n.d.) to create a gold standard dataset through manual annotation of 100 representative images related to COVID-19 and neurodegeneration. To systematically compare GPT outputs with these curated annotations, we established a single, unified BioBERT-based semantic matching pipeline that was applied consistently across all analyses. In this pipeline, both CBM gold-standard triples and GPT-generated triples are first normalized and concatenated into full subject–predicate–object strings. These are embedded using the BioBERT model (“dmis-lab/biobert-base-cased-v1.1”) (Bhasuran, 2022; J. Lee et al., 2020), and cosine similarity scores are computed between every GPT triple and every CBM triple from the same image.

To assign matches, we use the Hungarian algorithm, which finds the optimal one-to-one mapping between predicted and gold triples that maximizes overall similarity. The Hungarian algorithm (a.k.a. the Kuhn–Munkres algorithm) solves the linear assignment problem: given two sets and a cost (or similarity) matrix between their elements, it finds the optimal one-to-one matching that minimizes total cost (or maximizes overall similarity) (Kuhn, 1955; Shih & Parthasarathy, 2012). A predicted triple is considered a match if its similarity to the assigned gold triple meets or exceeds a predefined threshold, the exact value of which was evaluated further. This ensures that matches reflect strong semantic alignment while avoiding duplicate assignments.

Triples meeting or exceeding the similarity threshold were counted as true positives (TP), unmatched GPT triples as false positives (FP), and unmatched CBM triples as false negatives (FN). The aggregated results provided the overall baseline performance of the GPT-based extraction method, serving as a reference point for subsequent optimization steps.

For each evaluation, counts of TP, FP and FN are aggregated across the entire dataset. From these counts, precision, recall, and F1 scores are calculated (Blair, 1979; Chinchor, 1992; Manning et al., 2008). This baseline approach is applied unchanged for threshold optimization, prompt engineering comparison, and hyperparameter tuning, ensuring that all downstream results are directly comparable.

#### 2.3.3. Full-text triple comparison

In addition to figure-based extraction, we also evaluated GPT-4o for knowledge extraction directly from full-text biomedical articles. For this purpose, we implemented a dedicated pipeline that ingests cleaned article text, applies deterministic GPT prompting (S2 Table), and outputs structured semantic triples. The same schema as in the figure-based extraction was used, ensuring consistent subject–predicate–object formatting across sources.

To maintain comparability, we applied the same BioBERT-based semantic evaluation framework described above, using cosine similarity scoring and the Hungarian algorithm to align GPT-extracted full-text triples with CBM-curated gold-standard triples. As with figures, matched triples above the similarity threshold were considered TPs, unmatched GPT triples as FPs, and unmatched CBM triples as FNs.

### 2.4. Model selection and rationale

#### 2.4.1. Threshold evaluation

To assess the quality of GPT-extracted triples, we compared them against CBM-curated gold-standard triples subset using BioBERT embeddings and Hungarian alignment. Performance was tested across multiple similarity thresholds (0.70–0.90), and 0.85 was selected as the optimal cutoff, balancing precision and recall (see Supplementary Methods for details).

#### 2.4.2. Prompt engineering and hyperparameter tuning

We tested three prompting strategies: a free-form prompt, a fixed-predicate prompt (based on BEL (Biological Expression Language) relations), and a balanced hybrid (Hoyt et al., 2018; Slater, 2014). The free-form prompt (Prompt 1) yielded the best overall F1-score and was chosen for all downstream analyses. Full prompt texts and performance metrics are provided in Tables S3–S4.

Using the optimal prompt configuration, we systematically evaluated the effect of GPT-4o decoding parameters on triple extraction. Specifically, we varied temperature and top_p across five settings (1.0, 0.75, 0.5, 0.25, 0.0) and generated triples for a CBM-processed image evaluation subset. The resulting outputs were compared against the CBM gold standard using the evaluation framework described above, and precision, recall, and F1 metrics were calculated to quantify the impact of decoding parameters (for details see Supplementary Methods).

#### 2.4.3. Triple categorization

To enable downstream interpretation, all triples were mapped into six biological domains: (i) Viral Entry and Neuroinvasion, (ii) Immune and Inflammatory Response, (iii) Neurodegenerative Mechanisms, (iv) Vascular Effects, (v) Psychological and Neurological Symptoms, and (vi) Systemic Cross-Organ Effects. Categorization was performed using BERT-based embeddings and MeSH-derived dictionaries (see Supplementary Methods). Two complementary classification modes were applied: a Subject–Object mode (for comparison with CBM data) and a Pathophysiological Process (PP) mode (for the full GPT corpus).

## 3. Results

### 3.1. Evaluation of GPT-4o in relevant image assessment

To evaluate reliability, we manually reviewed a random sample of 500 image URLs. The comparison yielded accuracy = 0.90, precision = 0.48, recall = 0.96. These results indicate that GPT-4o effectively captured nearly all relevant images (high recall) but admitted some irrelevant ones (moderate precision), reflecting a deliberate design bias toward inclusivity.

### 3.2. Evaluation of threshold, prompt, and hyperparameter settings

We systematically evaluated GPT-4o extraction quality under three optimization strategies: similarity threshold selection, prompt engineering, and decoding hyperparameters (Fig. 2).

**Fig. 2.**
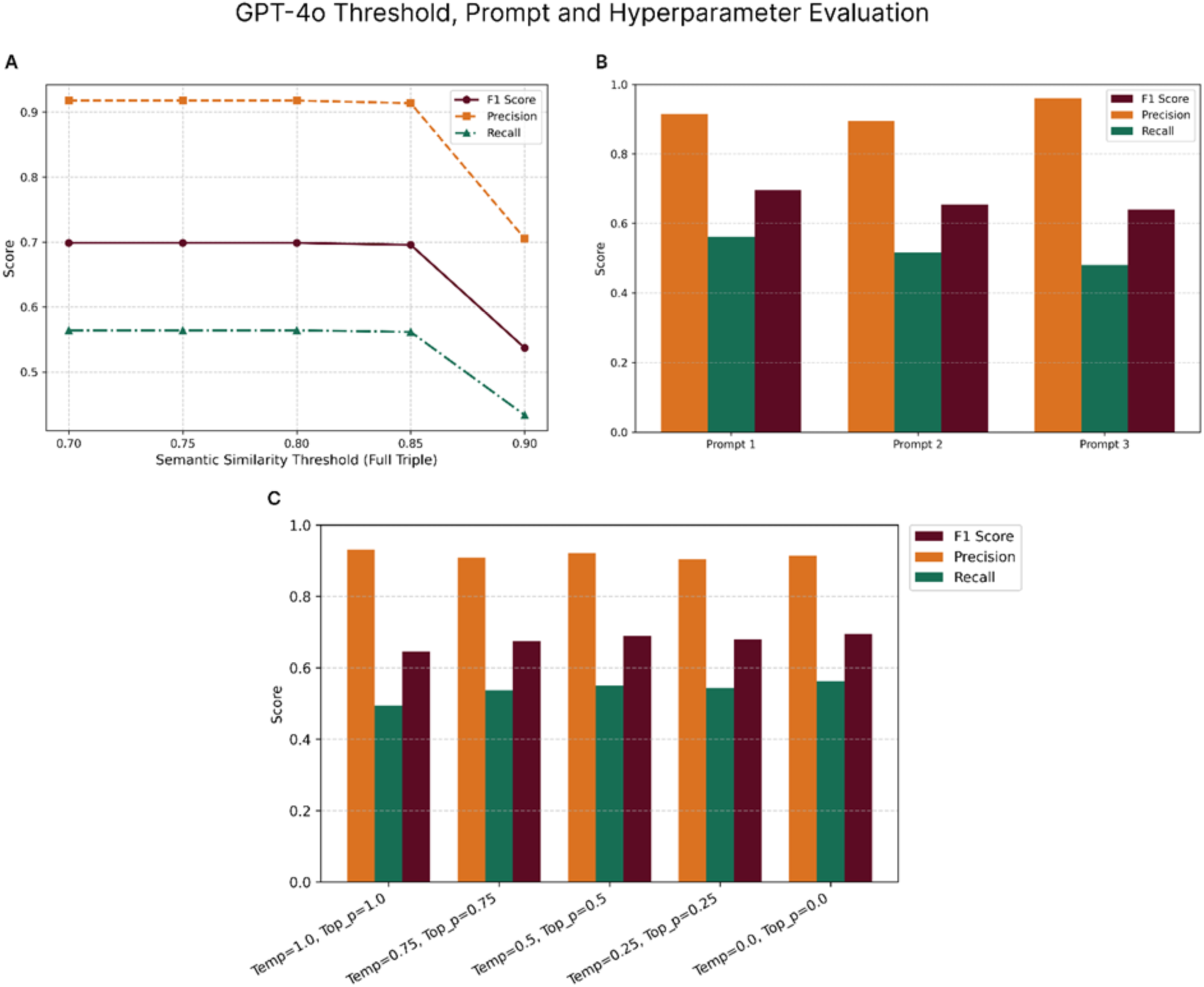
Evaluation of threshold, prompt, and hyperparameter settings for GPT-4o triple extraction. (A) Threshold optimization for semantic similarity alignment. (B) Comparison of prompt configurations. (C) Effect of decoding parameters (temperature, top_p). Precision, recall, and F1-scores are shown in all panels.

First, we tested semantic similarity thresholds for aligning GPT triples with CBM gold-standard triples using BioBERT embeddings and Hungarian alignment. F1-scores were relatively stable between 0.70–0.85 but declined sharply at 0.90, while recall dropped substantially at the highest cutoff. A threshold of 0.85 was therefore selected as the optimal balance between precision and recall (S5 Table).

Second, we compared three prompt configurations differing in constraint level. The free-form extraction (Prompt 1) achieved the best overall performance, balancing precision (0.91) and recall (0.56) for the highest F1-score (0.70). The constrained prompt (Prompt 2) underperformed due to reduced recall, while the hybrid (Prompt 3) achieved very high precision (0.96) at the cost of recall. On average, however, five out of 50 images could not be processed by GPT-4o due to refusals, likely triggered by the model’s sensitivity filters for medical content (*OpenAI API documentation*, 2024; Wang et al., n.d.). These refusals were inconsistent across runs and were not caused by access issues, as all CBM-provided images were independently hosted and verified. Full prompt texts and detailed evaluation metrics are provided in Supplementary Tables S3 and S6.

Finally, we assessed the impact of decoding parameters (temperature, top_p). While precision remained high across all configurations, recall and F1-scores varied. The deterministic setting (temperature = 0.0, top_p = 0.0) yielded the most consistent and reliable outputs, supporting its use for downstream KG construction. Full results are summarized in S7 Table.

### 3.3. Triple categorization

To facilitate mechanistic interpretation, extracted triples were categorized into six biological domains of COVID-19 pathophysiology. For comparison with the CBM gold standard, categorization was based on Subject and Object terms only, ensuring methodological consistency. As shown in Fig. 3A, both datasets emphasize Immune & Inflammation and Cross-Organ Effects, followed by Viral Entry & Neuroinvasion, despite differences in absolute counts.

**Fig. 3.**
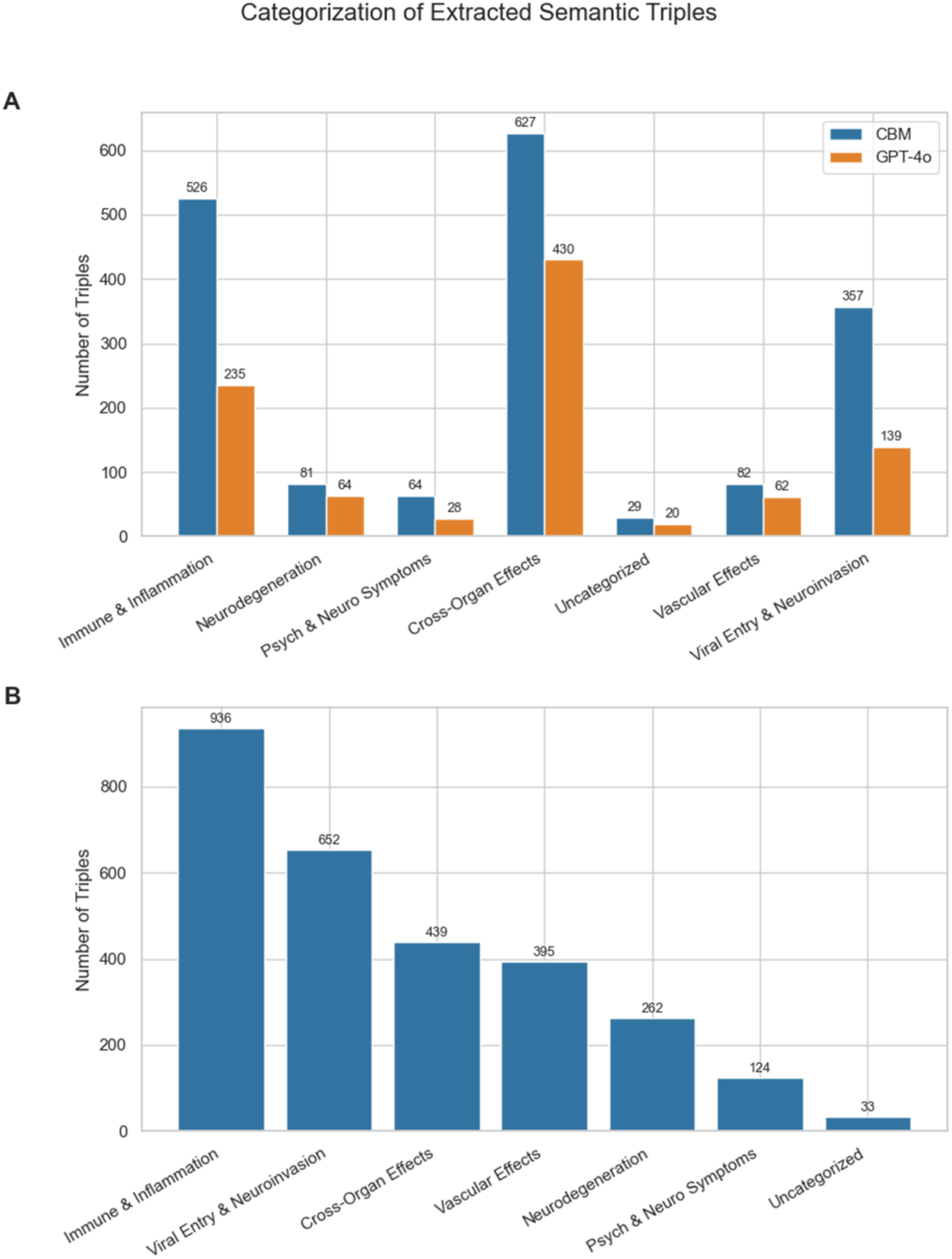
Categorization of extracted semantic triples. (A) Comparison between CBM and GPT-4o on the 93-image evaluation set, categorized via subject and object terms. (B) Full distribution of GPT-4o triples (289 images), categorized using the PP field.

For the full GPT-4o corpus (289 images), categorization was performed using Sentence-BERT similarity against MeSH descriptors, with fallback GPT classification for ambiguous cases. The resulting distribution (Fig. 3B) again highlighted immune signaling, neuroinflammation, and viral entry as dominant mechanisms. These results reinforce the biological coherence and topical relevance of the GPT-4o–extracted triples, further validating their potential for enriching KGs focused on COVID-19–NDD comorbidity.

### 3.4. KG visualization and analysis

To enable graph-based exploration and analysis, we utilized the Neo4j graph database platform (Ian Robinson, et al., 2015). Triples were ingested via a custom Python pipeline, where subject and object nodes were heuristically assigned to predefined biological categories based on a combination of fuzzy string matching, lexical cues, and ontology lookups. This initial step leveraged manually curated keyword lists (e.g., for "cell", "protein", or "biological process") and ontology APIs such as the Disease Ontology, Gene Ontology, HGNC, and MESH (Braschi et al., 2019; Lipscomb, 2000; Schriml et al., 2019; The Gene Ontology Consortium, 2021). Entities were matched to known terms using approximate string similarity (e.g., via RapidFuzz (Bachmann, 2020/2025)), structural heuristics (e.g., gene name patterns), and semantic hints from their descriptions. This allowed us to automatically infer node labels such as Gene, Disease, or Cell_Phenotype in a domain-aware yet scalable manner.

However, given the inherent ambiguity of biological terms and the potential for misclassification in heuristic methods, we implemented a second refinement stage using GPT-4. After importing the pre-labeled nodes into Neo4j, we queried GPT-4 with each entity and its assigned label, prompting it to verify or correct the classification based on a fixed controlled vocabulary of nine high-level biomedical categories (e.g., Protein, Phenotype, Chemical). GPT-4’s outputs were used to reassign more accurate labels, correcting misclassifications and standardizing edge cases. At the end, the nodes that could not be assigned to one of the relevant types and were too general or of type None and Unknown were removed from analysis. This hybrid approach—combining rule-based heuristics with LLM-driven validation—yielded improved accuracy and interpretability of graph annotations. A representative visualization of the finalized, harmonized KG is shown in Fig. 4.

**Fig. 4.**
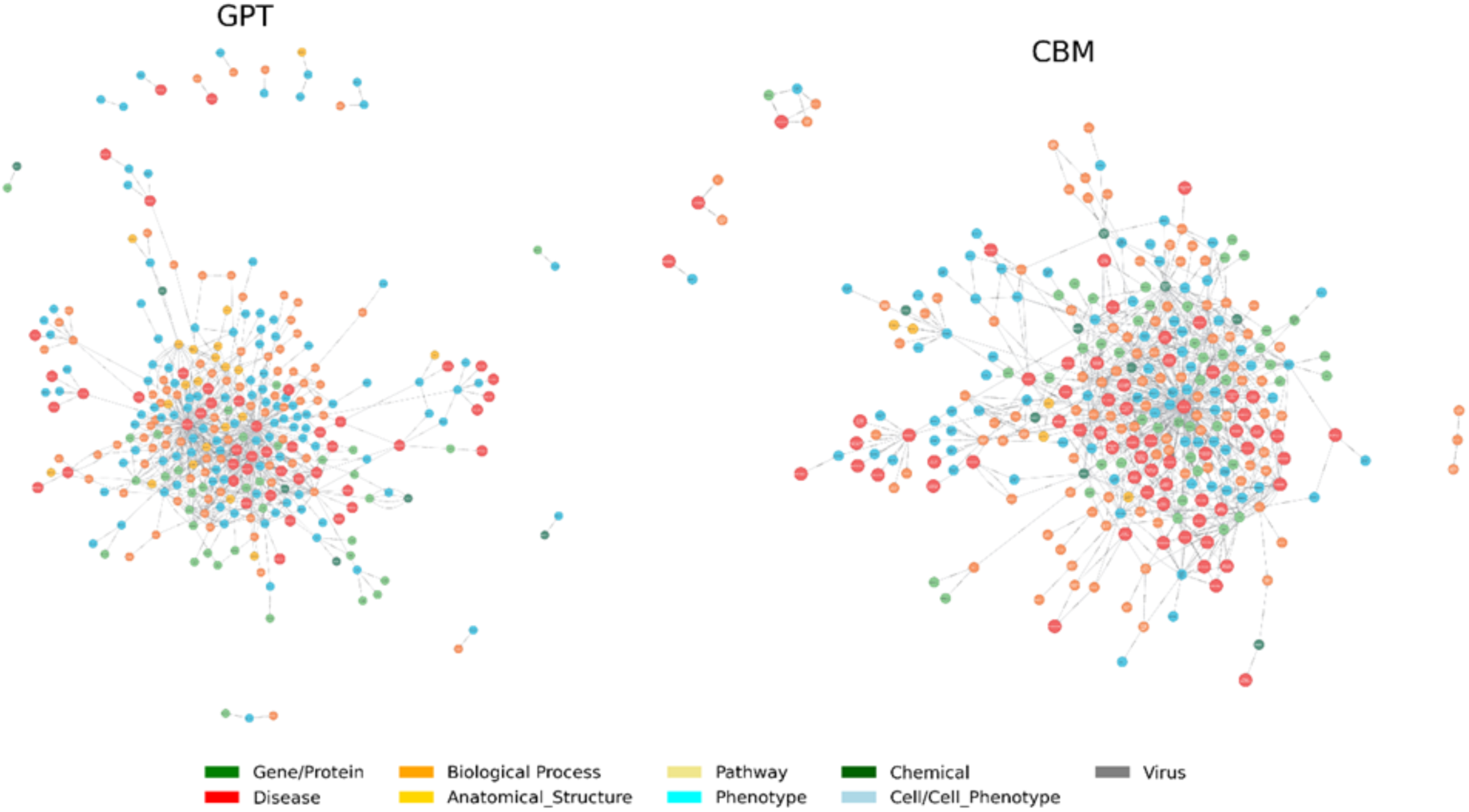
Visual comparison of KGs generated from GPT-4o and CBM triples. Triples extracted by GPT-4o (left) and CBM-curated gold standard (right) were imported into Neo4j for structural analysis. Each node represents a biomedical entity colored by neo4j label (e.g., Gene/Protein, Disease, Chemical). Edges represent semantic relationships between entities.

The manually curated and GPT-extracted KGs both aim to represent the complex web of SARS-CoV-2 biology, yet they differ substantially in scale, consistency, and biological depth (Fig. 5). The CBM network encompasses 861 nodes and 1,573 edges, yielding a density of 0.0048, whereas the GPT-4o graph is smaller — 791 nodes linked by 878 edges (density 0.0031). Visually, CBM forms a tightly interwoven core of processes, molecules and cell types, with only a handful of small, disconnected components. In contrast, GPT’s network is looser: its core is less dense, and it contains numerous small islands of nodes, reflecting both missed extractions and a proliferation of semantically overlapping labels. These topological differences are mirrored in the distribution of node and relationship types between the two networks (Fig. 6), with CBM emphasizing curated regulatory clarity and GPT-4o surfacing diverse mechanistic vocabulary. Both graphs identify *Biological_Process* as the single largest class—approximately 200 nodes in CBM versus 150 in GPT — but CBM thereafter drops steeply into *Gene*, *Chemical* and *Disease* categories, each with 40–75 nodes, before tapering into *Protein*, *Phenotype*, *Cell* and rarer types. GPT, by contrast, produces a far flatter spectrum: *Protein*, *Chemical*, *Disease*, *Cell* and *Anatomical_Structure* all appear at 25–30 nodes, with *Gene*, *Phenotype* and *Cell_Phenotype* lingering in the high tens and an almost negligible *Unspecified* category. CBM densely populated the network with the prototypical cytokines, interferons and chemokines you find in the literature. GPT, working from its broad statistical grasp, surfaces a more heterogeneous and under-sampled handful of genes— only one bona fide chemokine (CCL11), one inflammasome gene (NLRP3) and several non-canonical factors (e.g. MOF, ACSL4) (Fig. S1). A similar contrast appears in relationship-type usage. CBM deliberately normalizes every up-regulation or down-regulation statement into just two predicates—*INCREASE* (∼1,200 edges) and *DECREASE* (∼200 edges)—plus a handful of specialized interactions (e.g., *POSITIVECORRELATION*, *INFECT*). This concise ontology facilitates systematic analysis of regulatory cascades, such as the TNF–IL6 feedback loop driving vascular permeability in acute respiratory distress syndrome. GPT, on the other hand, retains a wider vocabulary—*CAUSES* (∼300), *LEADS_TO*(∼150), *INDUCES*, *TRIGGERS*, *AFFECTS*, *ACTIVATES*, *RESULTS_IN*, and more—scattering edges across fifteen distinct predicates. At the level of individual hubs, both graphs highlight the same biological components but with dramatically different connectivity. Across the protein hubs, CBM places the SARS-CoV-2 virus itself at roughly 70 connections, TNF at ∼20, the original SARS-CoV at ∼10 and both the spike glycoprotein and ACE2 receptor at about eight each. In contrast, GPT’s protein layer is much sparser: the SARS-CoV-2 spike glycoprotein and ACE2 each appear with only ∼6 edges, a generic *spike protein* node at ∼4, other viral components at ≤3. At the gene level, CBM is dominated by canonical cytokines and chemokines—IL-6 (26), IL-1β (18), IFN-γ and IL-10 (∼7), IFN-β, IL-2 and CXCL8 (4) and CXCL10 (3)—whereas GPT-4o surfaces only a handful of genes: CCL11 (3), NLRP3, LIP and NF-κB (2 each) plus a smattering of stress or epigenetic regulators (HPA, MOF, ATF3, ACSL4, SLC7A11, M1 at 1 each) (Del Valle et al., 2020). Likewise, CBM’s cell-phenotype layer is anchored by activated microglia and M1 astrocytes (7 edges apiece), but in GPT-4o the top cell phenotypes are activated microglia (6) and infected immune cells (5), with M1 astrocytes down at only 3. Finally, neither network elevates specialized processes such as synapse pruning into high-degree hubs.

**Fig. 5.**
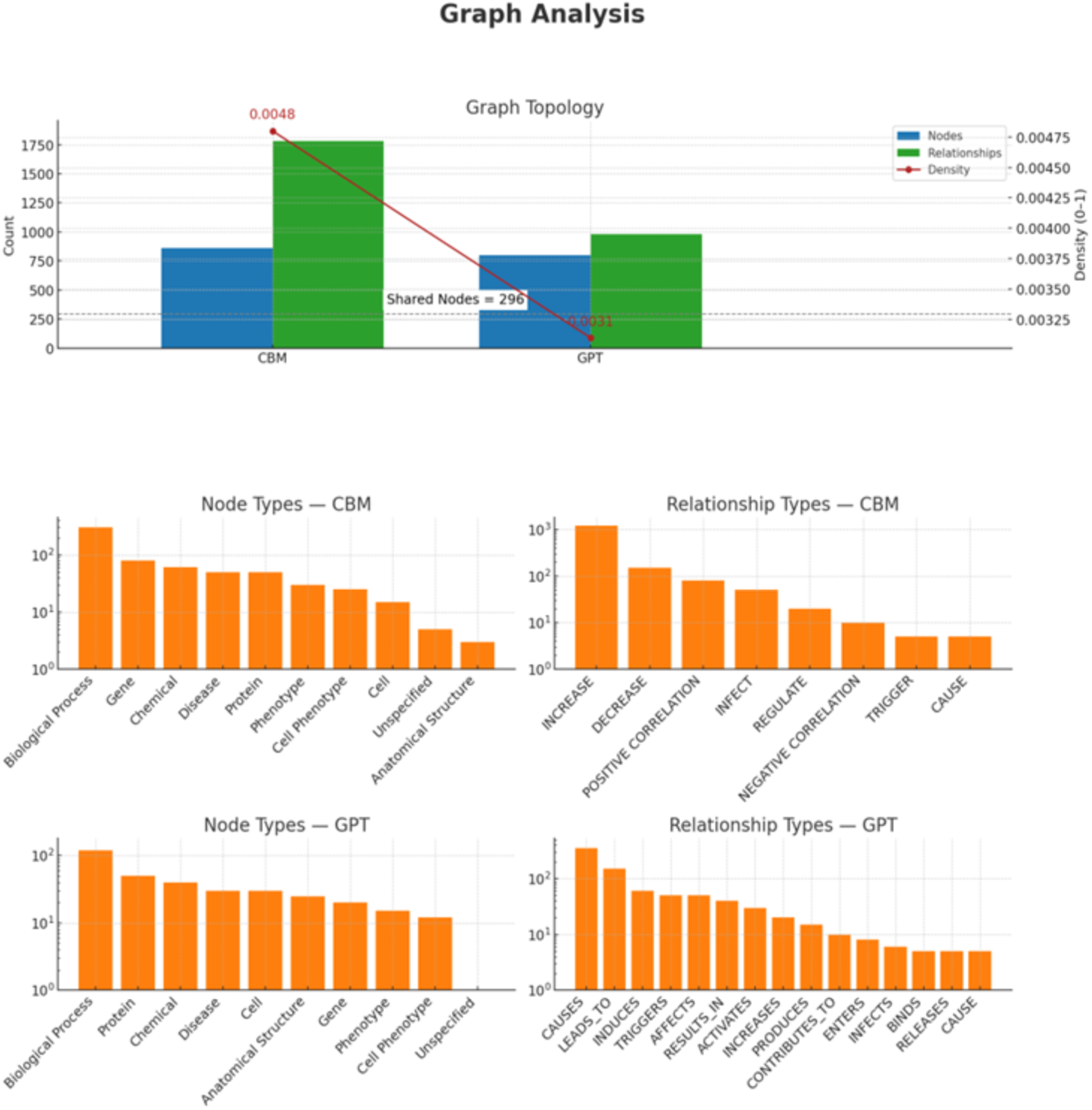
Comparison of extracted mechanisms from CBM and GPT-4o KGs. Side-by-side Neo4j-rendered layouts of KGs built from GPT-extracted and CBM-curated semantic triples. Nodes are color-coded by entity type, showing differences in graph density, connectivity, and semantic scope. Bar plots show the top 20 most frequently occurring mechanisms in each corpus (CBM top, GPT-4o bottom), excluding general categories (e.g., *COVID-19*) and missing annotations. CBM emphasizes high-level clinical conditions and phenotypes, while GPT-4o surfaces molecular and cellular processes.

**Fig. 6.**
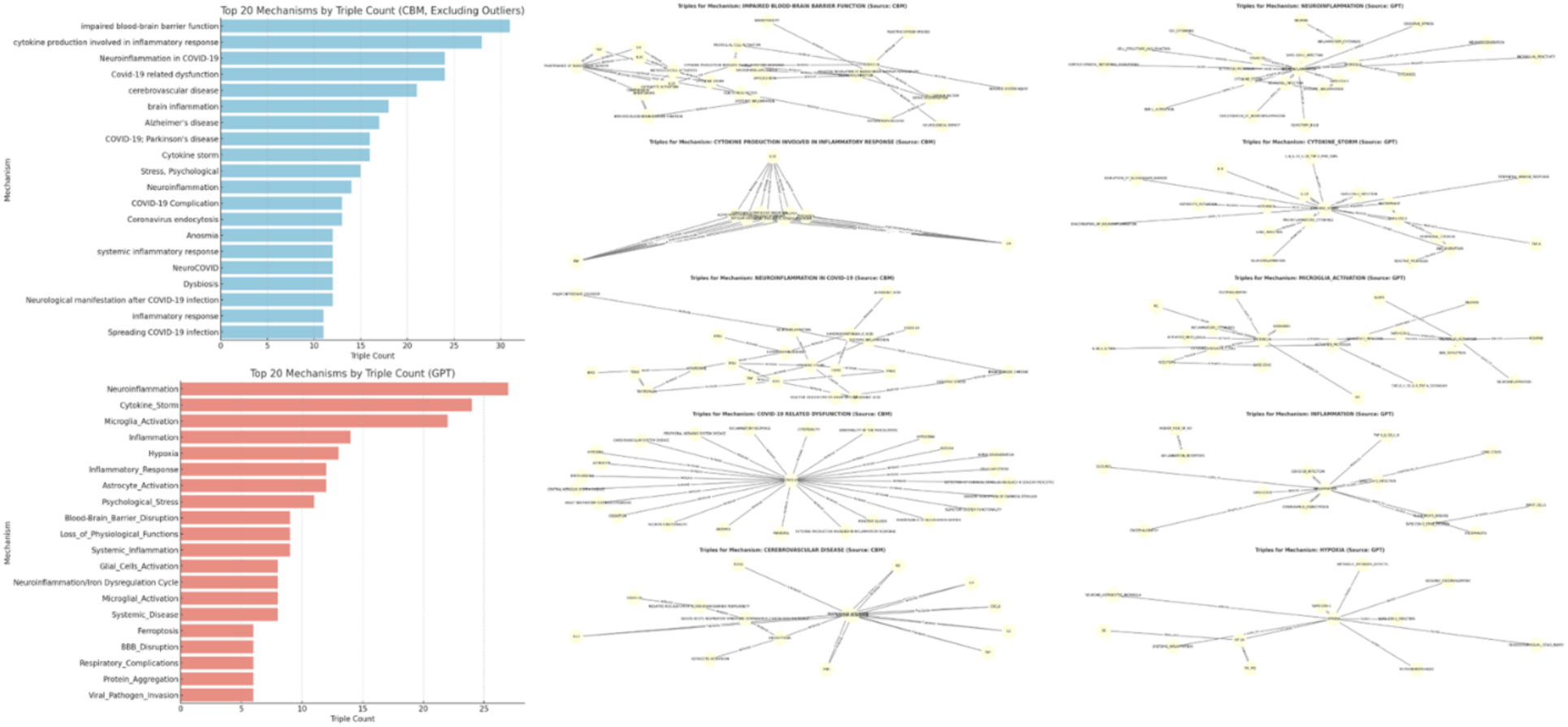
Comparative analysis of top mechanisms and network structures extracted from CBM and GPT-4o sources. (Left): Bar charts showing the top 20 mechanisms ranked by triple count from the CBM gold-standard corpus (top, blue) and GPT-4o-extracted triples (bottom, red). (Right): Example subgraphs illustrating the semantic triples associated with select high-ranking mechanisms from each source.

After excluding outliers—such as overly general categories (e.g., COVID-19) and entries with missing labels—analysis of the top 20 mechanisms reveals a strong convergence between manual curation and LLM-based extraction. Both approaches consistently identify neuroinflammation and blood–brain barrier (BBB) disruption as central drivers of COVID-19’s neurological manifestations and their intersection with NDD pathways. Nonetheless, each method emphasizes complementary aspects: manual curation tends to prioritize high-level clinical phenotypes (e.g., AD, PD, cerebrovascular disease, anosmia, systemic inflammatory response), whereas the LLM more frequently highlights underlying cellular and molecular processes (e.g., microglial and astrocyte activation, hypoxia, ferroptosis, protein aggregation, viral invasion) (Fig. 6, Fig. S1.).

While both sources underscore immune-mediated mechanisms such as neuroinflammation and cytokine storm, differences are evident in their relative ranking, emphasis, and mechanistic scope. CBM-derived subgraphs exhibit denser, more coherent structures, reflecting the selectivity of expert-driven annotation. In contrast, GPT-4o-derived subgraphs (Fig. 6, bottom right) capture a broader but more heterogeneous landscape, encompassing a wider range of mechanistic associations that are less tightly clustered.

#### 3.4.1. Performance of GPT-4o in visual understanding

To evaluate the accuracy of GPT-4o in extracting structured biological knowledge from biomedical figures, we conducted a systematic comparison against a gold standard of expert-curated annotations developed by biomedical specialists at CBM. This reference set comprised 93 context-agnostic images annotated with mechanistic triples related to COVID-19 and neurodegeneration, of which 7 could not be analyzed due to access restrictions or GPT-4o processing errors. We employed semantic similarity to assess the match between GPT-generated and gold-standard triples, as described in the Materials and methods section.

Table 2 summarizes the results of comparing GPT-4o-generated triples to the CBM gold standard using BioBERT full-triple embeddings and Hungarian algorithm alignment with a cosine similarity threshold of 0.85. The model achieved a precision of 0.917, indicating that the vast majority of predicted triples aligned semantically and biologically with the curated reference set. The F1 score of 0.675 reflects a balanced but imperfect performance, while recall remained moderate at 0.534, showing that some curated relationships were not recovered.

**Table 2.** Performance metrics for GPT-4o triple extraction compared to CBM Gold Standard.

| Metric | Value |
| --- | --- |
| True Positives (TP) | 897 |
| False Positives (FP) | 81 |
| False Negatives (FN) | 783 |
| Precision | 0.917 |
| Recall | 0.534 |
| F1 Score | 0.675 |
| Semantic Threshold | 0.85 |

Performance metrics are calculated globally across the full image set, used for CBM manual curation. The comparison was preformed using BioBERT-based full-triple embeddings and Hungarian alignment at a cosine similarity threshold of 0.85.

Despite this gap in recall, qualitative review revealed that many FPs captured plausible but unannotated mechanisms — for example, ferroptosis, glial activation, and transcriptional regulators (e.g., MOF, ACSL4) — which extend beyond the current scope of the gold standard. GPT-4o also identified intermediate processes such as oxidative stress, astrocyte polarization, and barrier permeability, adding mechanistic detail to the curated dataset’s endpoint-focused annotations. These findings suggest that, although coverage is incomplete, GPT-4o’s positive predictions are highly reliable and biologically relevant.

Fig. 7A–B presents the triple classification breakdown and aggregated evaluation metrics, while Fig. 7C shows the distribution of cosine similarity scores for all GPT–CBM triple pairs. The histogram demonstrates that most predictions — including many non-matches — are semantically close to their nearest gold standard counterpart, with similarity values clustering between 0.80 and 1.00. This pattern highlights GPT-4o’s capacity to produce biologically meaningful triples even when they do not meet strict match criteria.

**Fig. 7.**
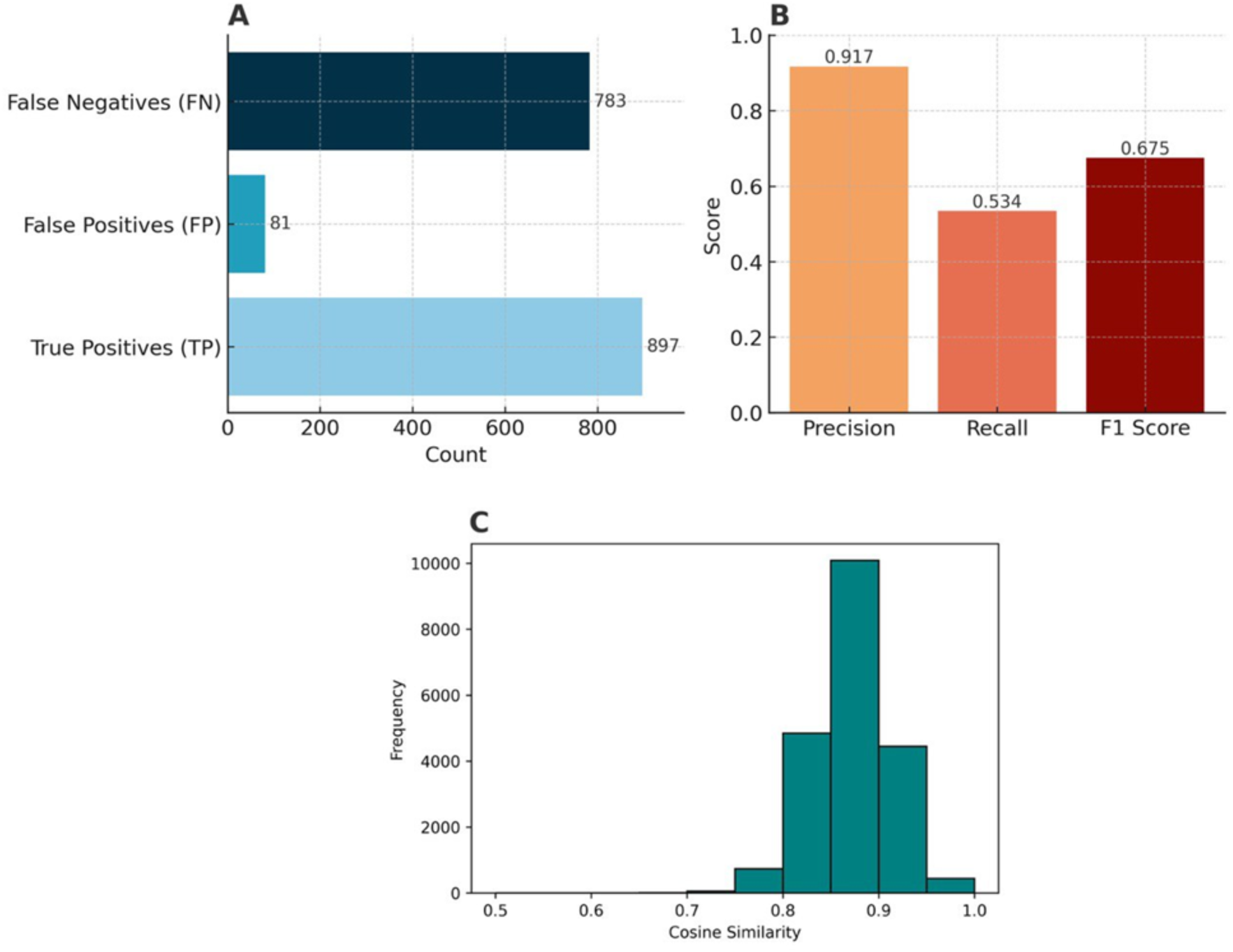
Performance of GPT-4o triple extraction compared to CBM Gold Standard using BioBERT semantic matching. (A) Triple classification breakdown for GPT-4o predictions vs. CBM gold standard. (B) Precision, recall, and F1 score computed at a cosine similarity threshold of 0.85. (C) Distribution of cosine similarity scores for all GPT–CBM triple pairs, showing clustering in the high-similarity range (0.80–1.00).

To complement the aggregate metrics in Table 2, we examined representative matched and unmatched triple pairs from the evaluation, as shown in Table 3. These examples illustrate how GPT-4o’s BioBERT-based full-triple similarity scoring and Hungarian one-to-one alignment operate in practice under a cosine similarity threshold of 0.85. The qualitative cases reveal how the system balances semantic overlap with biological specificity, and how the threshold and alignment constraints contribute to correct matches, correct rejections, and classification errors.

**Table 3.** Examples of matched and unmatched triple pairs during evaluation.

| Image | GPT-4o Triple | CBM Triple | Cosine Similarity | System Decision | Interpretation |
| --- | --- | --- | --- | --- | --- |
| image_5 | lung injury causes hypoxia | lung injury increases hypoxia | 0.991 | Match | **True Match** – The GPT and CBM triples express the same mechanistic relationship with near-identical entity and predicate wording, resulting in both semantic and biological equivalence. |
| image_78 | excitotoxicity contributes to neurodegenerative and neuropsychiatric diseases | ace2 decreases infections | 0.663 | No Match | **True Mismatch** – The two triples describe unrelated biological processes, and the low similarity score correctly leads to a non-match classification. |
| image_1 | sars cov 2 infection causes hypoxia | covid 19 increases cytotoxic t cell infiltration | 0.870 | Match | **False Positive** – Although the similarity score is near the threshold, the triples differ biologically: one describes respiratory consequences, the other immune infiltration. The match is semantically misleading. |
| image_1 | hypoxia leads to ischemia | covid 19 increases cxcl10 | 0.841 | No Match | **False Negative** – The triples are biologically related via hypoxia-driven inflammatory signaling, but fall just below the similarity threshold, resulting in a missed match. |

Examples of matched and unmatched GPT-4o–CBM triple pairs at a semantic similarity threshold of 0.85, illustrating representative cases of true matches, true mismatches, FPs, and FNs. Each example includes cosine similarity, system decision, and an interpretation of the classification outcome in terms of semantic and biological alignment.

Overall, the examples in Table 3 demonstrate that GPT-4o’s conservative similarity threshold favors precision over recall, effectively filtering unrelated relationships but occasionally excluding biologically valid ones. Many near-threshold non-matches reflect conceptually related processes, suggesting that modest threshold adjustments or secondary biological reasoning modules could recover additional relevant triples without substantially compromising precision.

#### 3.4.2. Comparative analysis with text mining

To evaluate the added value of image-derived information relative to text, we constructed three KGs: CBM (expert-curated from figures), GPT (images), and GPT-fulltext (Figs. 8–9, Table 4). All were generated using identical prompting strategies and harmonized parameters to ensure comparability.

**Fig. 8.**
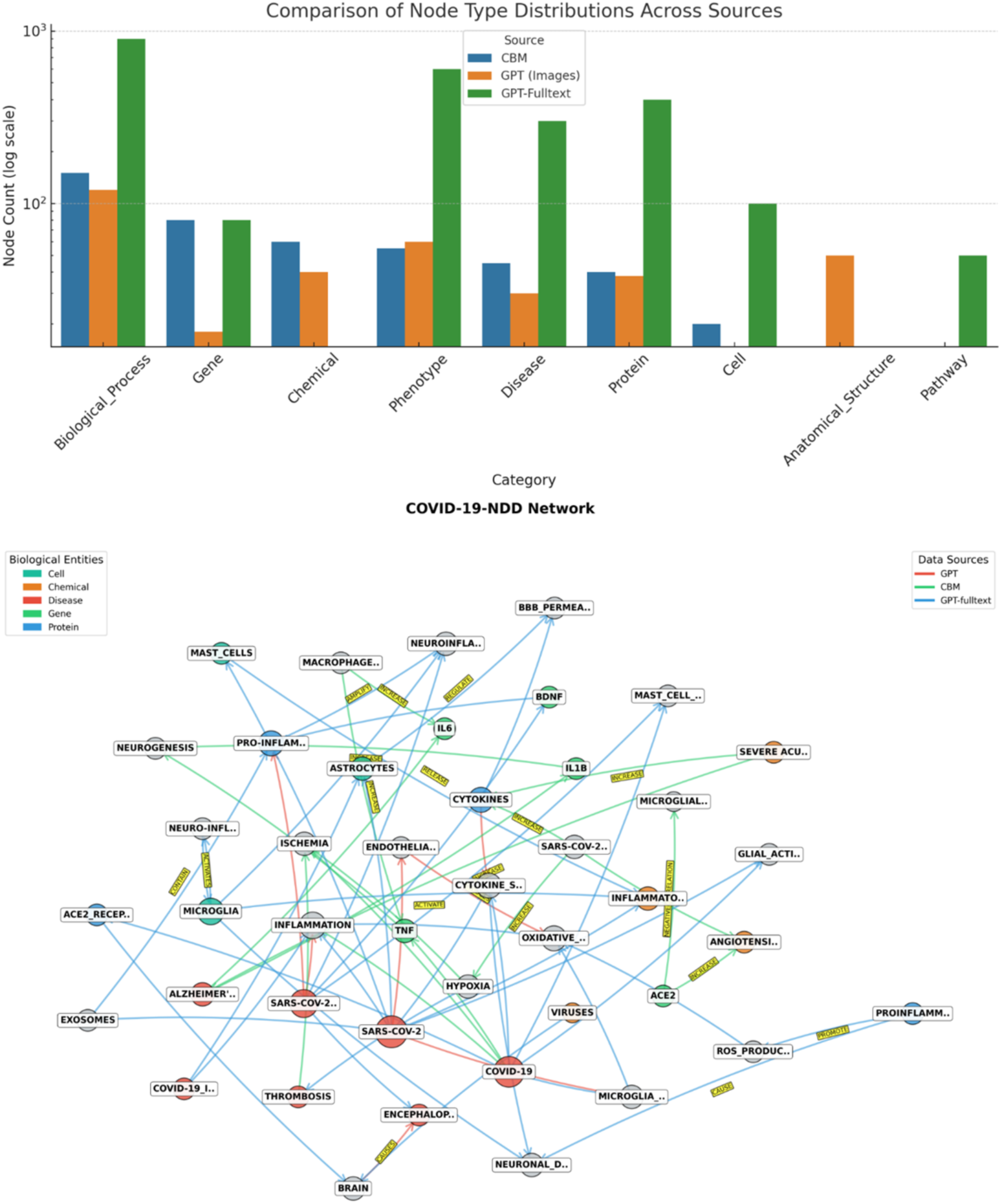
Comparative network analysis highlights distinct signatures of knowledge extraction across multi-modal sources. Comparative network analysis of CBM (green), GPT-images (red), and GPT-fulltext (blue). CBM emphasizes dense, molecularly precise interactions, GPT-images capture figure-derived causal scaffolds, and GPT-fulltext provides broad but diffuse coverage. The bar chart shows differences in entity type distributions.

**Fig. 9.**
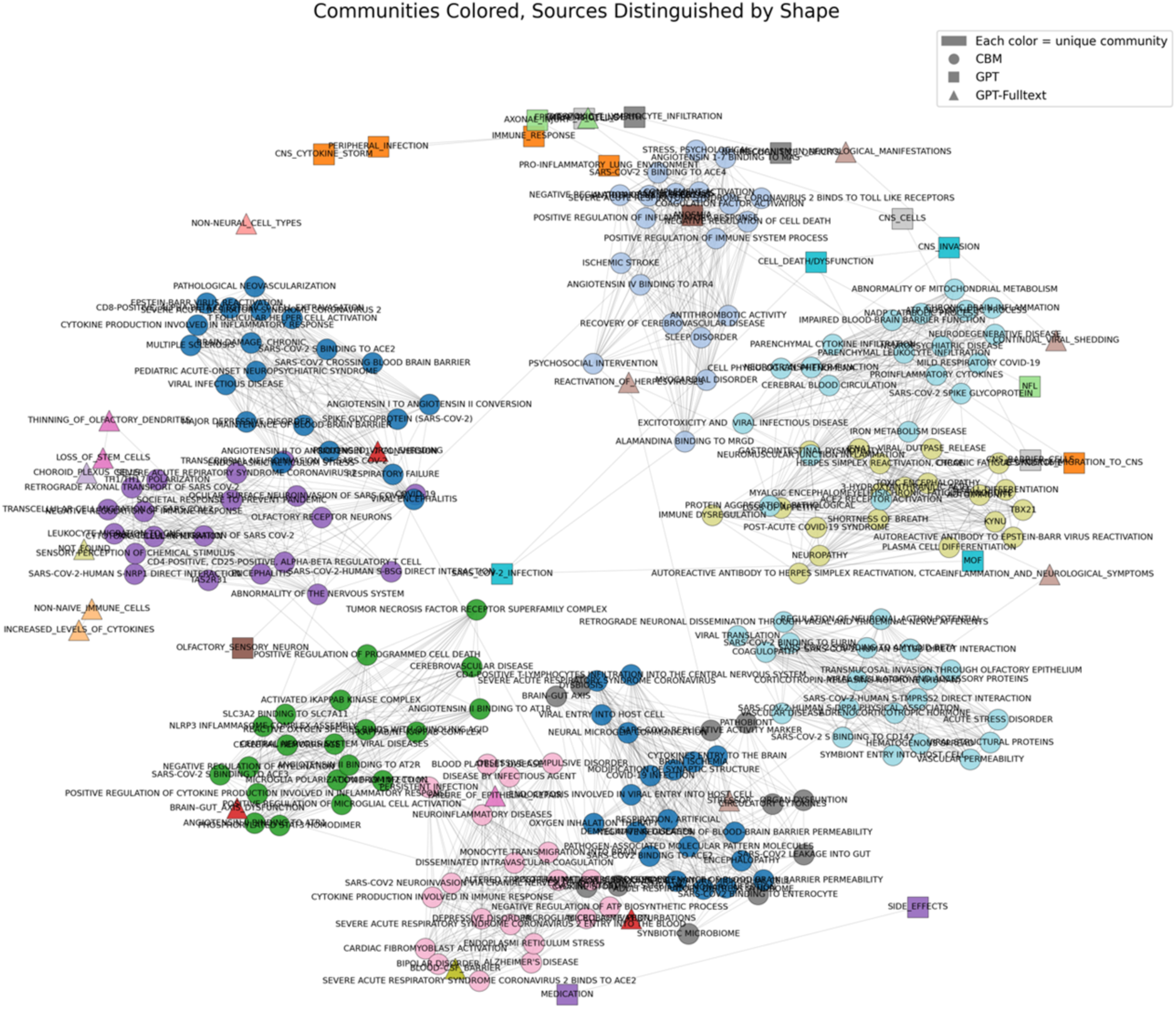
Combined community network across CBM, GPT, and GPT-Fulltext sources. CBM forms dense, biologically coherent clusters linking COVID-19 with neurodegenerative phenotypes, while GPT-derived networks are sparser and more fragmented, highlighting exploratory associations.

**Table 4.** Key characteristics and biases of KGs derived from manual curation, image-based extraction, and full-text mining.

| Source | Nodes | Edges | Density | Top Node Types (by count) | Relationship Style | Focus & Likely Bias |
| --- | --- | --- | --- | --- | --- | --- |
| **CBM (manual curation)** | 861 | 1573 | 0.0042 | Biological Process, Gene, Chemical | INCREASE / DECREASE, correlations | Expert-driven, molecular specificity (e.g., IL-6, TNF, ROS); denser, curated local subgraphs; focus on mechanistic depth. |
| **GPT**<br>**(images)** | 791 | 878 | 0.0028 | Biological<br>Process,<br>Phenotype,<br>Anatomical<br>Structure | Causal<br>predicates<br>(CAUSES,<br>LEADS-TO,<br>INDUCES) | Figure-derived, causal<br>scaffolding across<br>scales (cytokine storm<br>→ BBB disruption →<br>neurological<br>outcomes); emphasizes<br>clinical/phenotypic<br>outcomes over<br>molecular detail. |
| **GPT-**<br>**fulltext** | 6397 | 7352 | 0.0004 | Biological<br>Process,<br>Phenotype,<br>Protein, Disease | Causal<br>predicates<br>(CAUSES,<br>ACTIVATES,<br>INDUCES) | Literature-wide,<br>breadth and<br>redundancy; reinforces<br>well-known<br>associations (ACE2,<br>IL-6, cytokine storm)<br>but can overweight<br>generic<br>virus/inflammation<br>links; broadest<br>coverage, lowest<br>density. |

CBM yields dense, molecularly precise graphs; GPT (images) captures causal scaffolds linking systemic inflammation to neurological outcomes; GPT-fulltext provides the broadest but sparsest graphs, reflecting literature-wide redundancy.

The graphs differed substantially in scale and connectivity. CBM yielded 861 nodes and 1,573 edges (density 0.0042), GPT (images) contained 791 nodes and 878 edges (density 0.0028), while GPT-fulltext produced the broadest network with 6,397 nodes and 7,352 edges but the lowest density (0.0004). These differences highlight methodological trade-offs: manual curation creates small but densely interconnected networks emphasizing mechanistic precision; automated figure extraction produces mid-scale causal scaffolds aligned with diagrammatic logic; and full-text mining yields expansive but diffuse networks reflecting the narrative style of biomedical writing.

Despite these contrasts, all three approaches converged on neuroinflammation, blood–brain barrier (BBB) disruption, and cytokine storm as central mechanisms linking COVID-19 and neurodegeneration. Cross-modal validation identified 84 high-confidence relationships, with especially strong agreement for COVID-19 → neuroinflammation, neuroinflammation → oxidative stress, and oxidative stress → brain damage, as well as clinical manifestations such as COVID-19 → anosmia, dysgeusia, and encephalopathy. BBB dysfunction pathways were also consistently recovered across modalities, including SARS-CoV-2 → BBB disruption and cytokine storm → BBB disruption. Importantly, one relationship—COVID-19 → inflammation—was validated across all three sources, marking it as the highest-confidence mechanistic link.

Each modality contributed complementary strengths. CBM emphasized molecular biomarkers and inflammatory mediators (e.g., IL-6, TNF, ROS), reflecting expert annotation of experimental detail. GPT (images) captured figure-level causal scaffolds such as systemic inflammation → cytokine storm and SARS-CoV-2 → IL-6, bridging systemic and clinical outcomes. GPT-fulltext expanded coverage to broader processes such as renin–angiotensin imbalance, oxidative stress, and microglial activation, albeit at the cost of lower density. Together, these perspectives underscore the complementarity of curated, visual, and textual data.

Community detection further revealed structural contrasts (Fig. 9). CBM produced coherent clusters embedding COVID-19 within immune and neurological phenotypes, highlighting mechanistic overlaps such as post-viral fatigue and pediatric neuropsychiatric disorders. In contrast, GPT-derived networks were smaller and fragmented, capturing scattered associations that serve primarily as hypothesis generators. While less interpretable, these exploratory links suggest potential novel connections warranting further validation.

Overall, integration of curated, image-based, and text-derived graphs yields a multi-scale KG that spans molecular mechanisms, biological processes, and clinical outcomes. This synthesis reinforces confidence in convergent mechanisms while highlighting modality-specific contributions: CBM for depth, GPT-images for causal scaffolds, and GPT-fulltext for breadth. Additional insights, illustrative triples, source-exclusive lists, node-level vocabularies, and extended community details are provided in Supplementary Results.

### 3.5. Biological insights from the multimodal KG

#### 3.5.1. Shared inflammatory hubs between COVID-19 and neurodegeneration

We analyzed nodes that were connected within two steps to both COVID-19 and NDD reference nodes, thereby identifying shared hubs. This analysis confirmed a convergence on inflammatory mediators (Fig. 10; Fig. S2). The curated dataset (CBM) strongly emphasized the canonical cytokine triad IL-6, TNF, IL-1β, which are well-known drivers of systemic and neuroinflammation (Del Valle et al., 2020). GPT extractions (from figures and full text) not only captured this triad but also broadened the inflammatory network to include chemokines CXCL10, CXCL12, and CCL2, and in the case of full-text mining, IL-17, pointing to adaptive immune involvement. This indicates that manual curation reference nodes the KG in canonical inflammatory axes, while GPT methods introduce under-sampled chemokine and antibody-related processes, enriching the representation of post-COVID neuroinflammatory signatures.

**Fig. 10.**
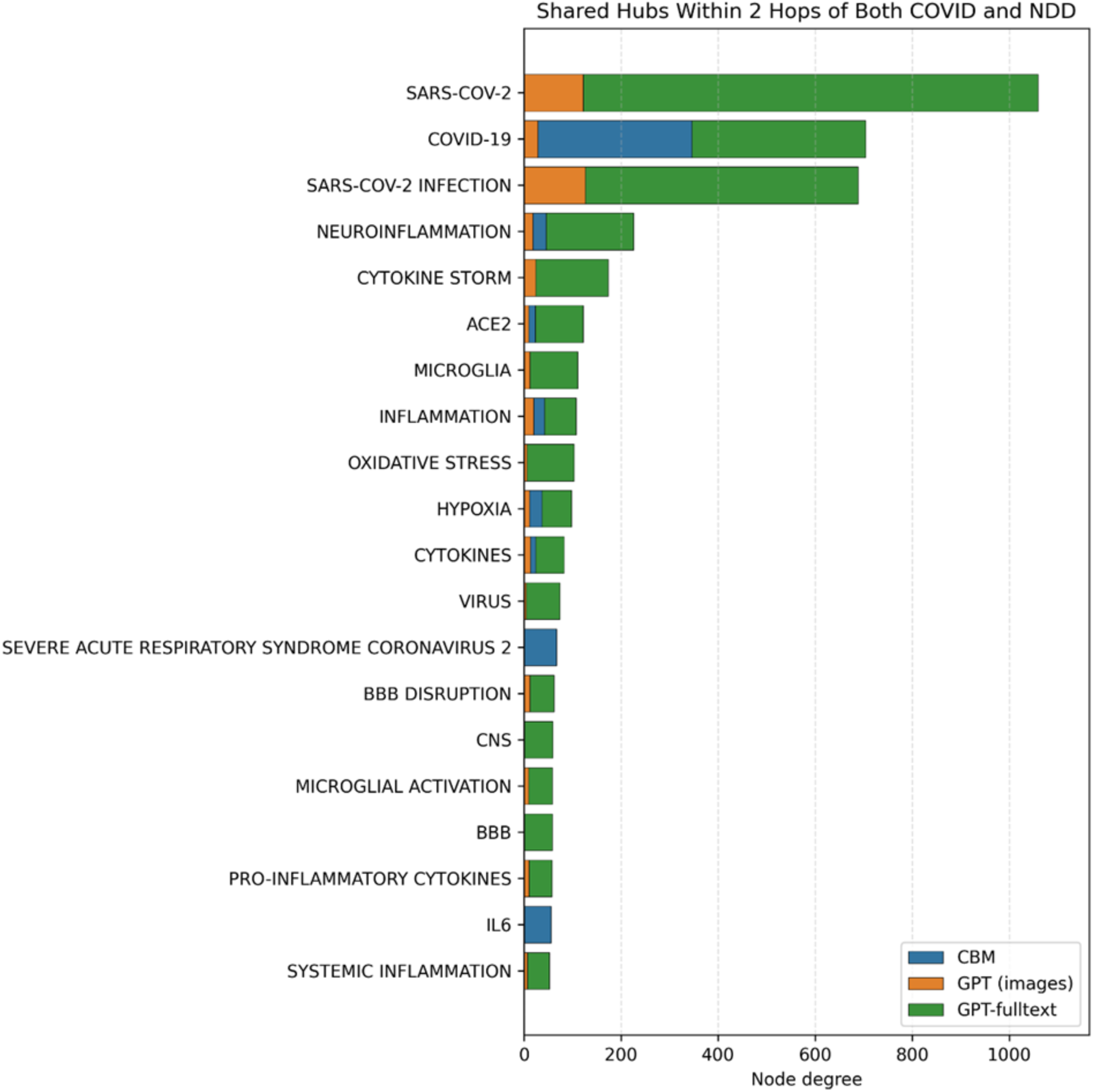
Shared hubs connecting COVID-19 and neurodegeneration. Top 20 nodes within two hops of both COVID and NDD reference nodes ranked by degree, split by source (GPT, CBM, GPT-fulltext). Canonical cytokines dominate, while GPT enriches chemokine and antibody-related hubs.

#### 3.5.2. COVID-19–induced BBB disruption and glial activation

Shortest-path analysis between COVID-19 and blood–brain barrier nodes revealed two mediator classes (Fig. S3). CBM annotations concentrated on neuroinflammation as the primary bridge, reflecting a canonical pathway. GPT extractions, however, highlighted systemic contributors such as cytokine storm, hypotension, and macrophage activity, suggesting alternative vascular or immune routes for barrier compromise.

Neighborhood analysis of astrocyte activation reinforced this complementarity (Fig. S4). CBM concentrated on classical mediators (IL-6, TNF, IL-1β) and closely related processes such as cytokine production and microglial activation. GPT added exploratory links, including oxidative stress, autoantibodies, and diverse chemokines, pointing to possible novel modulators of glial reactivity. Together, these results show that curated data reliably capture established neuroinflammatory routes, while GPT enriches the landscape with systemic and less conventional mechanisms, including processes like ferroptosis surfaced elsewhere in the GPT-figure extractions.

#### 3.5.3. Therapeutic bridge candidates highlight minocycline

Applying a majority-beneficial filter (down-modulation of COVID-side inflammatory/viral/BBB processes and neuro-side pathological outcomes, supported by ≥2 sources) identified minocycline (Garrido-Mesa et al., 2013) as the top therapeutic bridge candidate (Table 5).

**Table 5.**
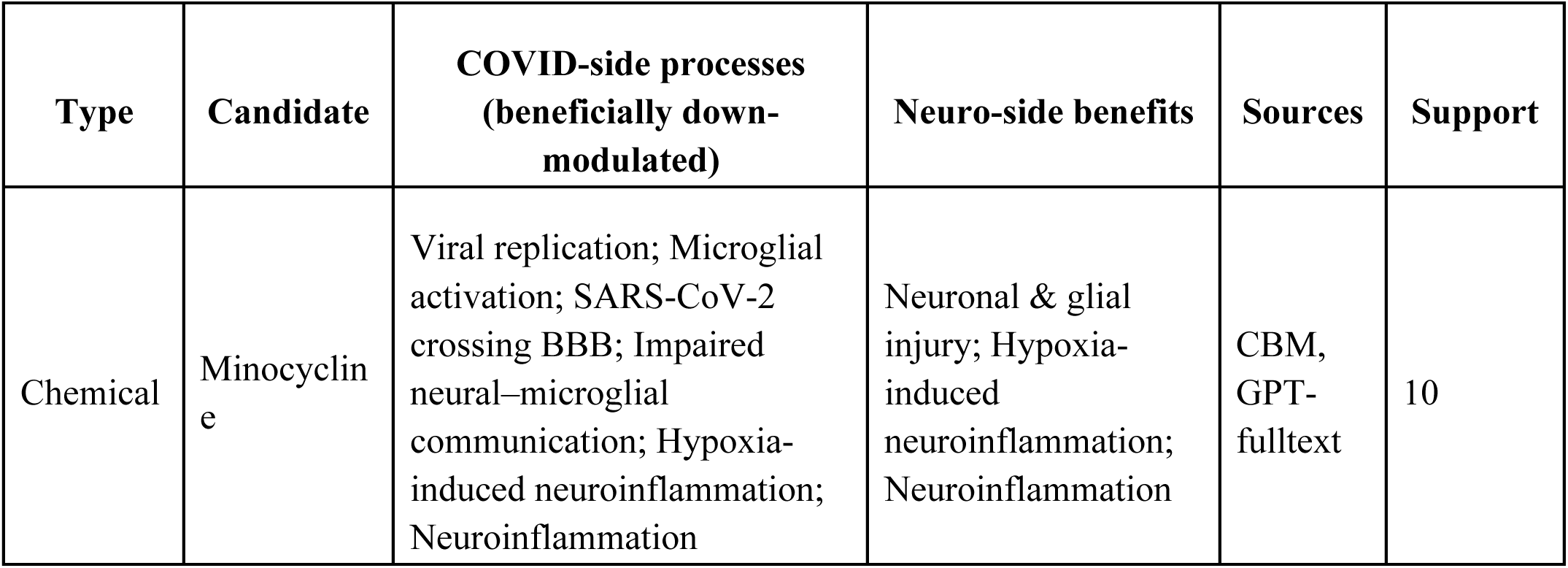
Therapeutic targets and repurposing opportunities.

| Type | Candidate | COVID-side processes<br>(beneficially down-modulated) | Neuro-side benefits | Sources | Support |
| --- | --- | --- | --- | --- | --- |
| Chemical | Minocycline | Viral replication; Microglial activation; SARS-CoV-2 crossing BBB; Impaired neural–microglial communication; Hypoxia-induced neuroinflammation; Neuroinflammation | Neuronal & glial injury; Hypoxia-induced neuroinflammation; Neuroinflammation | CBM, GPT-fulltext | 10 |

Therapeutic bridge analysis identified minocycline as the most consistent repurposing candidate, linking COVID-19–related inflammatory/viral/BBB processes with neuroprotective outcomes across modalities.

CBM curation annotated minocycline as decreasing microglial activation, viral replication, and SARS-CoV-2 crossing of the BBB. GPT-fulltext extractions connected minocycline to reduced neuroinflammation, hypoxia-induced neuroinflammation, and neuronal/glial injury, often via beneficial predicates such as ALLEVIATES, COUNTERACTS, PREVENTS (Fig. 11). Notably,

**Fig. 11.**
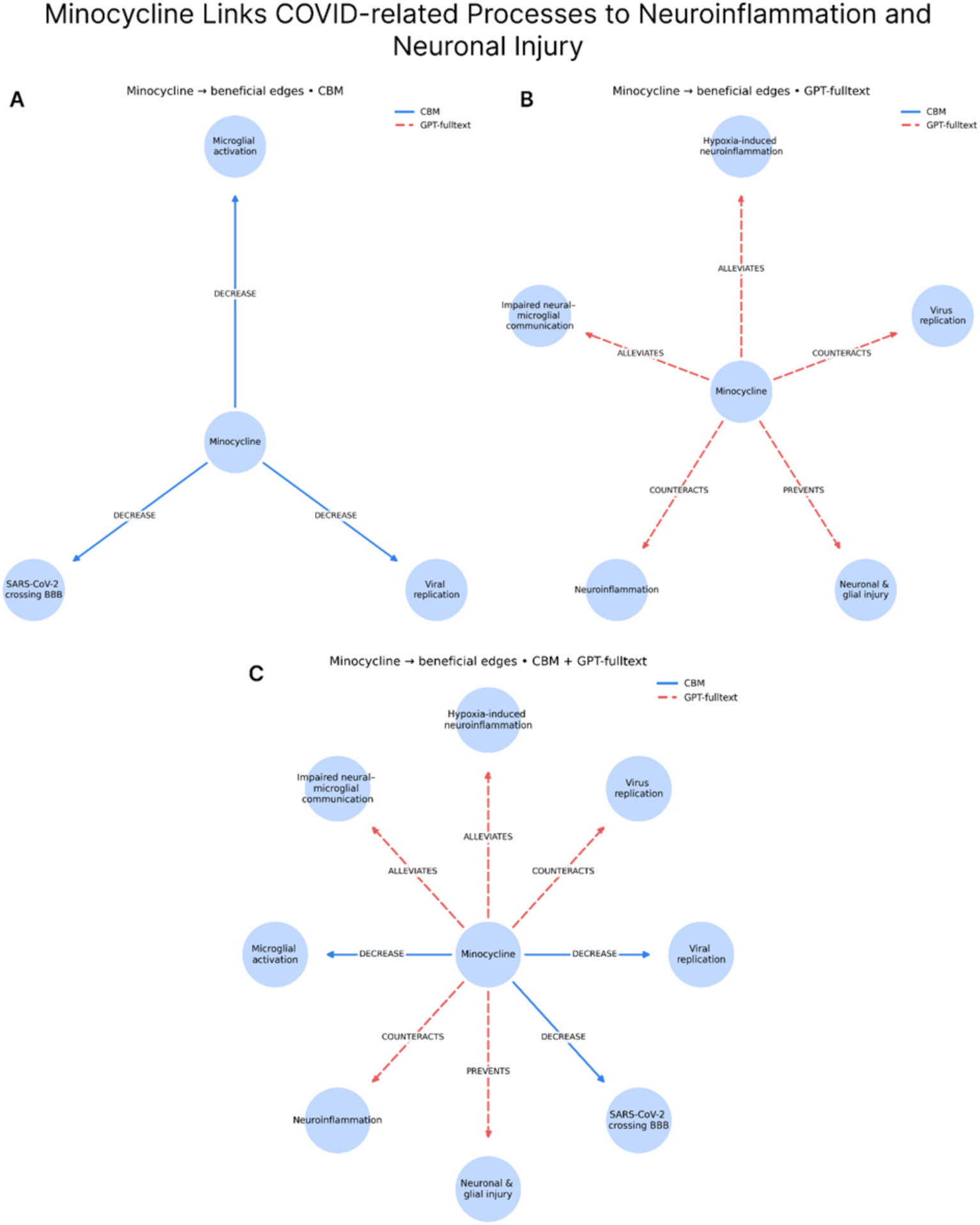
Minocycline links COVID-related processes to neuroinflammation and neuronal injury. (A) GPT-fulltext: connections to neuroinflammatory outcomes via ALLEVIATES, COUNTERACTS, PREVENTS. (B) CBM: decreases microglial activation, viral replication, and SARS-CoV-2 BBB crossing. (C) Combined view: complementary evidence across sources.

GPT-from-figures did not capture minocycline, highlighting source complementarity: figures enrich novel mechanistic processes (e.g., ferroptosis), whereas curated and textual sources capture pharmacological evidence.

This integration underscores minocycline’s potential as a repurposing candidate at the COVID–NDD interface, supported by its dual role in dampening viral/inflammatory cascades and mitigating neuronal injury.

## 4. Discussion

Understanding the mechanistic links between COVID-19 and NDDs remains an open challenge, largely because biomedical knowledge is fragmented across multiple modalities—narrative text, curated databases, and scientific figures. This work introduced a systematic framework that directly addresses this gap by extracting biological relationships from graphical biomedical figures using GPT-4o, a state-of-the-art multimodal language model, and comparing them with relationships obtained from manual curation (CBM) and full-text literature mining (Acosta et al., 2022). To our knowledge, this is one of the first studies to evaluate the complementarity of image-based, text-based, and curated approaches in constructing KGs of disease mechanisms (Chen et al., 2023).

A key observation from our analysis is that figures often contain mechanistic information that is more explicit and fine-grained than what can be recovered from text alone. For instance, GPT (images) captured relationships such as SARS-CoV-2 activates astroglia and hypoxia causes oligodendroglial injury, while CBM provided precise molecular assertions such as IL-6 increases brain inflammation and Angiotensin II increases ROS. These causal chains are rarely verbalized so directly in the full-text corpus, where they are typically collapsed into broader associations such as COVID-19 leads to neuroinflammation or COVID-19 causes BBB disruption (Islamaj Doğan et al., 2019). Likewise, figures surfaced underexplored or emerging hypotheses—ferroptosis results in neuronal death, viral protein induces protein aggregation—that were absent or only vaguely referenced in the text (Qiu et al., 2024; Sinha et al., 2021). These examples underscore how figures capture visual causal scaffolds linking molecular events, cellular responses, and systemic outcomes, thereby complementing the broader but often less mechanistically connected associations extracted from literature mining. Manual curation (CBM) provided a compact but highly precise set of triples, with a focus on canonical immune mediators (e.g., IL6 increases brain inflammation) and directional relations (INCREASE, DECREASE). GPT-fulltext extraction generated the broadest network, recovering literature-wide associations such as SARS-CoV-2 infection exacerbates multiple sclerosis symptoms and reinforcing hubs around ACE2, IL-6, TNF, and cytokine storm. The triangulation of these three modalities consistently converged on neuroinflammation and blood–brain barrier (BBB) disruption as central axes linking viral infection to neurological injury (P. Huang et al., 2023), while highlighting complementary emphases: molecular specificity from CBM, causal topology from images, and breadth and redundancy from text (M. H. Lee et al., 2022).

Several methodological innovations underpin this work. First, we systematically compared prompting strategies and hyperparameter choices for GPT-4o and found that deterministic decoding yielded the most coherent and reproducible extractions. Second, we developed a BioBERT-based similarity pipeline coupled with the Hungarian algorithm to align GPT-generated triples against curated gold standards. Third, we conducted comparative graph analyses to evaluate node distributions, edge densities, and hub structure across sources. These efforts provide not only novel results for COVID–NDD biology but also a reusable framework for evaluating multi-modal knowledge extraction pipelines (Haq et al., 2025; Warner et al., 2024).

The comparative analyses revealed clear source-specific emphases. CBM produced the smallest but most densely connected subgraph, enriched for proteins, chemicals, and molecular processes, and employing consistent directional predicates. GPT (images) captured mid-scale, figure-derived causal chains that often linked infection to systemic and clinical effects, but tended to generalize at the category level (e.g., cytokines rather than IL-6). GPT-fulltext yielded the largest and broadest graph, heavily skewed toward biological processes, diseases, and phenotypes, but relatively under-representing molecular entities. This skew reflects the narrative richness of the literature but also introduces redundancy and generic associations. Together, these findings demonstrate that each modality provides distinct insights—precision from curation, causal scaffolds from images, and breadth from text—that can be harnessed synergistically.

At the same time, the work revealed important limitations and methodological challenges. One limitation stems from node harmonization: entities such as “IL-6,” “IL6,” and “Interleukin-6” were often treated as distinct, inflating uniqueness estimates. Ontology-driven alignment (e.g., MeSH, UniProt, UMLS) will be essential for reliable integration. Another limitation is predicate heterogeneity in GPT outputs, which included naturalistic but inconsistent expressions (“causes,” “cause,” “induces,” “triggers”), complicating graph consistency. Similarly, GPT outputs exhibited a bias toward processes and phenotypes, under-representing specific molecules compared to CBM. In the case of GPT (images), ignoring figure legends and surrounding text limited contextual grounding, leading to ambiguous nodes such as “viral protein.” Additionally, a subset of biomedical images could not be processed by GPT-4o due to sensitivity constraints, reducing the representativeness of figure-derived knowledge (Wang et al., n.d.).

From a methodological perspective, several strategies could further improve performance. Few-shot prompting, in which curated example triples guide GPT during inference, may reduce relation variability and improve semantic alignment (Lewis et al., 2021). Retrieval-augmented generation (RAG) could inject domain-specific vocabularies or ontology definitions, enhancing both predicate and entity normalization. Multimodal fusion, incorporating figure captions and article context alongside images, would provide richer disambiguation and improve grounding. Finally, semi-automated curation pipelines—where GPT proposes candidate triples and human curators validate and refine outputs—offer a practical compromise between scalability and accuracy (Gao et al., 2024; Regino & dos Reis, 2025).

Beyond methodological advances, our integrated analysis also yielded concrete biological insights. The three modalities consistently converged on IL-6, TNF, and IL-1β as dominant shared inflammatory hubs linking COVID-19 to neurodegeneration, while GPT extractions broadened the landscape to include chemokines such as CXCL10, CXCL12, and CCL2, and adaptive signals like IL-17. Shortest-path and neighborhood analyses highlighted neuroinflammation as the canonical bridge to BBB disruption (Heneka et al., 2020; Mayer & Fischer, 2025), with GPT contributing complementary systemic routes including cytokine storm (Etter et al., 2022), hypotension, and oxidative stress. Finally, therapeutic filtering identified minocycline as a multi-source repurposing candidate that simultaneously attenuates viral/immune processes and neuroinflammatory injury. This observation is in line with ongoing clinical efforts, as a randomized controlled trial has already evaluated minocycline in COVID-19 patients (Esmaeilzadeh et al., 2023). Together, these findings underscore both convergence on well-established mechanisms and the potential for multimodal extraction to surface novel, translationally relevant hypotheses.

Biologically, the convergence of all three modalities on immune dysregulation, vascular dysfunction, and glial activation reinforces growing evidence for a convergent pathophysiological axis linking viral infection to neurodegeneration. The repeated emergence of neuroinflammation, cytokine signaling, endothelial barrier disruption, and microglial/astrocytic activation across modalities underscores their robustness as mechanistic hallmarks of COVID-19 neuropathology (Henstridge et al., 2019). At the same time, modality-specific signals highlight complementary contributions: CBM reference nodes canonical mediators such as IL-6 and TNF; GPT-images capture underrepresented processes such as ferroptosis, hypoxia, or oligodendroglial injury; and GPT-fulltext reinforces broad comorbid associations (e.g., multiple sclerosis, PD). Importantly, GPT-4o surfaced emerging actors (e.g., MOF, ACSL4) and processes (oxidative stress, epigenetic regulation) that are underexplored in curated resources, providing fertile ground for hypothesis generation and future validation.

Looking forward, this framework points to several avenues for future research. Ontology-based harmonization will enhance integration across modalities, while hybrid pipelines that combine curated precision, figure-derived causal logic, and literature-wide breadth will yield richer, more balanced graphs. Extending the approach to other biomedical domains—oncology, immunology, or rare diseases—will test generalizability and expand impact. Finally, coupling extracted KGs with experimental or clinical datasets offers a path toward closed-loop systems for hypothesis generation, validation, and refinement.

In summary, this study establishes the feasibility and value of multi-modal biomedical knowledge extraction. Manual curation, GPT (images), and GPT (full text) each capture distinctive but complementary perspectives, and their integration yields a more comprehensive and mechanistically informative representation of COVID-19 and NDD mechanisms. By demonstrating that figures encode latent causal knowledge that is otherwise missed, this work highlights the importance of including images in next-generation biomedical text mining and lays the groundwork for integrative, multimodal knowledge discovery in biomedicine.

## 5. Conclusions

This study presents a novel multimodal, AI-driven framework for systematically extracting mechanistic knowledge from biomedical figures and translating it into structured, interpretable KGs. Focusing on the intersection of COVID-19 and NDDs, our approach harnesses GPT-4o’s vision-language capabilities to transform complex visual content — often neglected in conventional literature mining — into a rich, machine-readable source of mechanistic insight.

Our analysis reveals that SARS-CoV-2 infection is consistently associated with key neurobiological processes such as blood-brain barrier disruption, microglial activation, oxidative stress, and neuroinflammation — features common to both acute viral pathology and chronic neurodegenerative conditions. The semantic triples extracted by GPT-4o not only aligned closely with expert-curated knowledge but also uncovered additional, underrecognized mechanisms (e.g., ferroptosis, synaptic dysfunction) that enhance the depth of mechanistic understanding and suggest new avenues for investigation.

Biologically, the multimodal KG highlighted canonical inflammatory mediators (IL-6, TNF, IL-1β) as convergent hubs linking COVID-19 to neurodegeneration, while GPT-based extractions broadened the network to include under-represented chemokines (CXCL10, CXCL12, CCL2) and adaptive signals such as IL-17. Complementary analysis of blood–brain barrier disruption and glial activation reinforced neuroinflammation as a core axis, with GPT surfacing systemic and vascular contributors including cytokine storm, hypotension, and oxidative stress (Cai et al., 2018). Finally, therapeutic filtering consistently identified minocycline as a cross-validated repurposing candidate, connecting viral/immune processes with neuroprotective outcomes across sources. These findings demonstrate that multimodal integration not only enriches methodological pipelines but also yields biologically and translationally relevant insights.

Methodologically, this work integrates visual search, prompt engineering, and ontology-informed classification into a coherent pipeline capable of high-throughput and biologically meaningful knowledge extraction. Comparative graph-based analyses show that LLM-generated KGs offer broader semantic coverage and mechanistic granularity, while expert curation retains clinical precision and terminological rigor — highlighting the complementary strengths of automated and manual approaches. This underscores the promise of hybrid, human-in-the-loop systems for enriching biomedical discovery.

By positioning scientific figures as primary, structured sources of knowledge, our framework bridges a critical gap between visual content and computational analysis in biomedical research. As multimodal models continue to mature, their integration into data mining and curation workflows holds transformative potential for accelerating hypothesis generation, pathway discovery, and translational applications across a wide range of complex disease domains.

## List of abbreviations

Abbreviation: Full Term
AD: Alzheimer’s Disease
AI: Artificial Intelligence
API: Application Programming Interface
BBB: Blood–Brain Barrier
BEL: Biological Expression Language
BioBERT: Biomedical Bidirectional Encoder Representations from Transformers
CBM: Causality Biomodels
CNS: Central Nervous System
FN: False Negative
FP: False Positive
TP: True Positive
TN: True Negative
GPT: Generative Pre-trained Transformer
GPT-4o: GPT-4 Omni
HGNC: HUGO Gene Nomenclature Committee
HTTP: Hypertext Transfer Protocol
KG: Knowledge Graph
LLM: Large Language Model
MeSH: Medical Subject Headings
NDD: Neurodegenerative Disease
NLP: Natural Language Processing
PD: Parkinson’s Disease
PMID: PubMed Identifier
PP: Pathophysiological Process
SPOKE: Scalable Precision Medicine Open Knowledge Engine
URL: Uniform Resource Locator
VLM: Vision Language Model

## Declarations

### Availability of data and materials

The data and source codes used in this study are available at: https://github.com/SCAI-BIO/covid-NDD-image-based-information-extraction.

## Declaration of competing interests

The authors declare that they have no known competing financial interests or personal relationships that could have appeared to influence the work reported in this paper.

## Funding sources

This research was supported by the Bonn-Aachen International Center for Information Technology (b-it) foundation, Bonn, Germany, and Fraunhofer Institute for Algorithms and Scientific Computing (SCAI). Additional financial support was provided through the COMMUTE project, which receives funding from the European Union under Grant Agreement No. 101136957.

## CRediT Author Statement

- **Elizaveta Popova**: Methodology, Software, Data Curation, Formal Analysis, Writing – Original Draft.
- **Marc Jacobs**: Conceptualization, Writing – Review & Editing.
- **Martin Hofmann-Apitius**: Conceptualization, Supervision, Project Administration, Funding Acquisition.
- **Negin Sadat Babaiha**: Conceptualization, Methodology, Software, Formal Analysis, Supervision, Writing – Original Draft, Writing – Review & Editing.

### Acknowledgements

We thank the entire COMMUTE consortium (for a list of partners see https://www.commute-project.eu/en/about.html) for valuable input and discussions.

### Declaration of generative AI in scientific writing

During manuscript preparation the authors used Grammarly and ChatGPT/GPT-4o solely to improve readability and language. After using these tools, the authors reviewed and edited the content and take full responsibility for the publication’s content.

---

**S1 Table.** Prompt used with GPT-4o for automated relevance assessment of biomedical images. This prompt instructs the model to classify each image as "Yes" (relevant), "No" (irrelevant), or "Uncertain" based on visual indicators of mechanistic links between COVID-19 and neurodegeneration. Criteria include content type, clarity, visual style, and the presence of both biomedical topics.

**S2 Table.** GPT-4o prompt used for automated extraction of semantic triples from full-text biomedical papers. The table lists the exact prompt template applied to each paragraph, instructing the model to identify PPs linking COVID-19 to neurodegeneration and to express them as normalized subject–predicate–object triples. The format enforces strict output structure (PPs and Triples fields) and specifies how to handle irrelevant or non-scientific text.

**S3 Table.** GPT-4o prompts used for prompt engineering tasks. GPT-4o prompts used for semantic triple extraction from biomedical images. Each prompt instructs the model to identify pathophysiological mechanisms related to COVID-19 and neurodegeneration, and to represent them in structured format. Prompts vary in constraint levels: Prompt_1 allows free-form predicates; Prompt_2 restricts predicates to a predefined BEL-derived list; Prompt_3 provides a controlled yet flexible set of example predicates. These variations support consistency testing and evaluation of extraction quality.

**S4 Table.** Full set of semantic triples extracted using GPT-4o prompts and manually curated for evaluation (Can be found in Supplementary_material_S4_Table.xlsx). This table contains all semantic triples used in the evaluation of GPT-4o prompt performance. Triples were extracted using three prompt variants (Prompt_1, Prompt_2, Prompt_3) and multiple hyperparameter settings for Prompt_1 (temperature and top_p ranging from 0.0 to 0.75). Additionally, the table includes manually curated gold-standard triples used as reference for similarity scoring. Each row indicates the image identifier, prompt version, hyperparameter setting (if applicable), subject, predicate, and object of the triple.

**S5 Table.** Representative semantic comparisons between GPT-predicted and CBM gold-standard triples (image_29) GPT triple: *inflammatory cytokines cause neuroinflammation* Threshold applied: 0.85 (BioBERT cosine similarity) Manually inspected GPT-predicted triples were matched against CBM ground truth triples using BioBERT similarity scoring. All shown examples exceeded the threshold of 0.85 and reflect biologically relevant mechanisms linking COVID-19, inflammation, and neurodevelopmental or neurodegenerative processes.

**S6 Table.** Performance of GPT-4o prompt variants against CBM gold standard triples. Precision, Recall, and F1-score are computed using BioBERT full-triple matching with a similarity threshold of 0.85. Prompt 1 achieved the best balance of precision and recall, resulting in the highest F1-score. Prompt 3 had the highest precision but lower recall, while Prompt 2 showed the weakest overall performance.

**S7 Table.** Impact of temperature and top_p on Prompt_1 extraction performance. Precision, recall, and F1 scores for GPT-4o triples generated with Prompt_1 under different decoding parameter settings, compared to the CBM gold standard using BioBERT full-triple matching (threshold = 0.85).

**Fig. S1.** Top 20 hub nodes by degree in CBM-curated and GPT-extracted KGs. Comparison of top 20 hub nodes (most connected entities) between CBM-curated and GPT-extracted semantic graphs. Horizontal bars represent node degree (number of incident edges). CBM nodes are shown in green; GPT nodes are shown in orange. The hub nodes highlight frequently occurring biomedical entities in each knowledge extraction modality.

**Fig. S2.** Cytokine/chemokine hubs near both COVID-19 and NDD. Subset restricted to cytokines/chemokines. CBM emphasizes IL-6, TNF, and IL-1β, while GPT adds CXCL10, CXCL12, CCL2, and IL-17, expanding the inflammatory repertoire.

**Fig. S3.** Top mediators on COVID↔BBB shortest paths. Stacked bar plot showing mediators linking COVID and BBB disruption. CBM strongly emphasizes neuroinflammation; GPT adds systemic factors (cytokine storm, hypotension, macrophages).

**Fig. S4.** Top neighbors of astrocyte activation. Top 15 neighboring nodes of astrocyte activation. CBM emphasizes IL-6, TNF, and IL-1β, while GPT contributes oxidative stress, autoantibodies, and chemokines.

Supporting materials are provided in **Supplementary_File_1**.

## Notes

https://github.com/SCAI-BIO/covid-NDD-image-based-information-extraction

### Competing Interest Statement

The authors have declared no competing interest.

