## Supplementary File for "Leveraging Multimodal Large Language Models to Extract Mechanistic Insights from Biomedical Visuals: A Case Study on COVID-19 and Neurodegenerative Diseases"

### **Supplementary methods**

#### **Image relevance assessment pipeline details**

##### **Deduplication and initial processing**

To enable scalable processing, we applied a perceptual hashing method (Manning et al., 2008; Samanta & Jain, 2021), which computes compact, content-aware hash codes that preserve visual similarity. This approach clusters near-duplicate diagrams, allowing us to retain only the highest-quality or highest-resolution image in each group. After deduplication, 6,319 unique URLs remained.

##### **Accessibility validation and Content-based filtering**

Each image URL was programmatically checked for availability and format validity using HTTP response codes and file type headers. Images with broken links, restricted access, or unsupported formats were removed, excluding 2,705 entries.

Remaining accessible images were classified using GPT-4o with a structured prompt (Table S1). Images were labeled as Relevant, Not Relevant, or Uncertain. Only those marked Relevant were retained for the next stage, while Uncertain images were flagged for manual review.

##### **Iterative refinement and manual quality review**

A second round of GPT-4o classification was applied to previously marked Relevant images to reduce borderline or ambiguous cases.

Finally, candidate images were manually inspected. Figures with clear biological labeling, pathway representations, or explicit depictions of molecular/cellular mechanisms linking SARS-CoV-2 and neurodegeneration were retained, producing a final set of 289 images.

#### **Model selection and rationale**

##### **Comparison threshold evaluation strategy**

To systematically assess the quality of GPT-generated triples, we conducted a semantic similarity-based evaluation against a curated gold standard. We randomly selected 50 images from the CBM image pool and compiled a subset of CBM triples corresponding to these images. For the same 50 images, we generated GPT triples using Prompt_1 with temperature = 0.0 and top_p = 0.0 to ensure deterministic output.

To quantify triple alignment, we encoded each full triple (subject–predicate–object) using BioBERT and computed cosine similarity between all GPT–CBM triple pairs per image. Rather than performing greedy or local matching, we applied the Hungarian algorithm to identify a globally optimal one-to-one mapping that maximizes overall similarity while enforcing a tunable threshold for match validity.

We evaluated performance over a range of similarity thresholds (0.70–0.90), computing performance metrics for each threshold. This process enabled principled threshold selection for downstream graph evaluation. A plot of evaluation metrics across thresholds was generated, and a detailed comparison log—including all matched and unmatched triples and similarity scores—was saved for each image.

##### **Prompt engineering strategy**

Effective extraction of structured biomedical knowledge from visual content depends not only on model capacity but also on prompt design. To optimize the consistency and quality of semantic triples extracted from biomedical figures, we evaluated three prompt configurations varying in constraint level and linguistic guidance. Each prompt was designed to influence how the model handles predicate formulation and triple structuring.

We tested the following prompt variants:

- Prompt_1 (Free-form extraction): A minimalistic zero-shot instruction that asked the model to extract semantic triples (subject–predicate–object) directly from images, without providing constraints on predicate vocabulary. Despite its simplicity, this prompt produced the most semantically consistent and relevant triples, and was thus selected as the baseline for downstream experiments..
- Prompt_2 (Fixed predicates): This configuration introduced an explicit list of standardized predicates derived from Biological Expression Language (BEL) (Hoyt et al., 2018; Slater, 2014). The aim was to enforce consistency and encourage the use of structured, ontology-aligned relationships across extracted triples. By constraining predicate vocabulary, this prompt was designed to align model outputs more closely with curated biomedical knowledge graph standards.
- Prompt_3 (Balanced approach): Combines Prompt 1’s free-form style with added formatting instructions and a small curated list of biologically intuitive predicates. This configuration aimed to reduce ambiguity without over-restricting model output.

To evaluate each prompt’s performance, we used a manually curated gold standard of semantic triples (CBM-extracted) created for the randomly picked subset of 50 biomedical figures. For each prompt, GPT-4o-generated triples were compared against the corresponding gold standard using BioBERT. The comparison was performed according to the Evaluation framework, described above.

From the matches between CBM and GPT-extracted triple, we computed standard evaluation metrics, aggregated across all images per prompt. Full prompt texts can be found in the S3 Table of the Supplementary_File_1, extracted triples for each configuration are listed in the S4 Table.

##### **Hyperparameter optimization**

Using the optimal prompt configuration, we systematically tuned GPT-4o's API parameters to balance diversity and precision in triple generation, specifically evaluating the effect of decoding hyperparameters — temperature and top_p — on the precision and consistency of semantic triple extraction (Nguyen et al., 2024; *OpenAI API documentation*, 2024; Renze & Guven, 2024; Windisch et al., 2024).

We tested five parameter configurations:

- temperature = 1.0, top_p = 1.0
- temperature = 0.75, top_p = 0.75
- temperature = 0.5, top_p = 0.5
- temperature = 0.25, top_p = 0.25
- temperature = 0.0, top_p = 0.0 (fully deterministic)

For each configuration, triples generated from the 50-image evaluation set were compared against the CBM gold standard as described in the Evaluation framework section. The evaluation produced precision, recall, and F1 score metrics for each configuration, enabling a quantitative comparison of the impact of decoding parameters on triple extraction performance.

#### **Automated triple categorization details**

To enable downstream mechanistic analysis, all extracted semantic triples were systematically categorized into one of six predefined biological domains reflecting COVID-19–related pathophysiology: (i) Viral Entry and Neuroinvasion, (ii) Immune and Inflammatory Response, (iii) Neurodegenerative Mechanisms, (iv) Vascular Effects, (v) Psychological and Neurological Symptoms, and (vi) Systemic Cross-Organ Effects.

Categorization was performed using a BERT-based embedding approach combined with MeSH-derived keyword dictionaries. First, all category-specific keywords and synonyms were compiled from MeSH descriptors and normalized (lowercasing, punctuation unification). Each triple was then represented as an input string and compared against the category embeddings using Sentence-BERT (all-MiniLM-L6-v2) (Reimers & Gurevych, 2019). Cosine similarity was used to assign triples to the category with the highest-scoring match, provided the score exceeded a threshold (0.5); otherwise, the triple was labeled as Uncategorized.

Two complementary classification modes were implemented to ensure comparability across datasets:

- Subject–Object mode (used for CBM–GPT comparison): triples were categorized based on normalized Subject and Object terms only, ensuring methodological alignment with CBM data that lack explicit process annotations.
- Pathophysiological Process (PP) mode (used for the full GPT-4o corpus): the PP field of each triple was embedded and classified directly. When triples could not be confidently assigned to any category (i.e., were labeled as Uncategorized by the BERT–MeSH similarity approach), a GPT-4o fallback classification was applied. In this step, the normalized triple text was provided to GPT-4o with the six predefined categories, and the model was prompted to select the most appropriate label.

### **Supplementary results**

#### **Comparative analysis with text mining details**

The full-text KG expanded beyond canonical molecular interactions to capture broader physiological and pathological cascades. Representative examples include:

- SARS-CoV-2 infection causes neurodegeneration … production of ROS and proinflammatory factors occurs in CNS parenchyma and cerebral vessels → triple: PRODUCTION_OF_ROS_AND_PROINFLAMMATORY_FACTORS OCCURS_IN CNS_PARENCHYMA_AND_CEREBRAL_VESSELS.
- COVID-19 and MS are linked to various immunological and neurological mechanisms … SARS-CoV-2 infection exacerbates multiple sclerosis symptoms → triple: SARS-COV-2_INFECTION EXACERBATES MULTIPLE_SCLEROSIS_SYMPTOMS.
- In contexts where microglia are already primed … exacerbated injury is possibly mediated by primed microglia → triple: EXACERBATED_INJURY IS_POSSIBLY_MEDIATED_BY PRIMED_MICROGLIA.
- The ACE2 receptor is found in neurons, glial cells, and other sites … SARS-CoV-2 uses neuropeptide signaling pathways → triple: SARS-COV-2 USES NEUROPEPTIDE_SIGNALING.

Such examples demonstrate how textual redundancy and narrative breadth increased the volume of extracted triples, including speculative or multi-step mechanisms.

Source-exclusive strengths were also observed. CBM uniquely captured cytokine–response associations such as IL-6 → inflammatory response and CXCL10 → inflammatory response, while GPT-fulltext identified complex anatomical and temporal cascades (e.g., SARS-CoV-2 → ependymal cell death → epithelial repair failure → dendrite thinning). GPT-images surfaced broader systemic interactions, such as SARS-CoV-2 infection → gut dysbiosis → dopamine decarboxylase and lockdown psychosocial stress → glial activation.

Node-level analysis reinforced these distinctions. CBM concentrated on canonical immune mediators and processes (IL-6, TNF, BBB disruption). GPT (images) emphasized phenotypic scaffolds (anosmia, astrocyte activation, cytokine storm). GPT-fulltext recapitulated literature-wide hubs (ACE2, TMPRSS2, oxidative stress, microglial priming), reflecting broad coverage. Relation vocabularies followed similar patterns: CBM normalized to increase/decrease relations, GPT-images favored causal predicates (causes, leads_to), and GPT-fulltext expressed the widest causal lexicon.

Community-level inspection (expanded from Fig. 11) showed CBM communities consistently clustering COVID-19 with immune and neurological phenotypes, including overlap with myalgic encephalomyelitis/chronic fatigue syndrome and gut–brain axis pathways. By contrast, GPT-derived communities were structurally sparse, often forming pairs or triplets of nodes without intermediate mechanistic processes. While insufficient for robust mechanistic inference, these weakly connected associations provide exploratory signals for hypothesis generation.

Taken together, the Supplementary analysis highlights how each modality contributes unique but complementary knowledge domains: curated images emphasize molecular biomarkers, GPT-images capture systemic scaffolds, and GPT-fulltext broadens the mechanistic landscape to include detailed cascades and underexplored anatomical structures.

### **Supplementary tables**

**S1 Table. Prompt used with GPT-4o for automated relevance assessment of biomedical images.**

**Prompt for the relevance assessment:**

*Image URL is given.*

Analyze this image and assign relevance to it: "Yes" for relevant images, "No" for irrelevant images, and "Uncertain" if the relevance cannot be assessed. Follow this classification:

Relevant:
Images which:
Сlearly demonstrate the relationship between Covid-19 and neurodegeneration (any neurological impacts).
Don't contain lots of text (no more than 500 characters).
Don't just depict a research outline.
Don't be just graphs or represent photos derived from scientific tools (microscopic, histological images) and data visualization.
Are likely to be cartoons drawn by article authors.

Irrelevant:
Unrelated images, for example just an image of a virus particle or a sick person.
Images where correct interpretation of the data is impossible.
Images which display insights into Covid-19 OR Neurodegeneration, if one is present and the other is missing.

Uncertain:
The relevance of the image cannot be confidently determined based on the visual content.

Your answer should contain only a final decision in the following format: No/Yes/Uncertain (without dots).

Don't write anything else!

**S2 Table. GPT-4o prompt used for automated extraction of semantic triples from full-text biomedical papers.**

**Prompt for the full-text triple extraction:**

*A text paragraph from a biomedical paper is given.*

Describe the following scientific paragraph from an article on comorbidity between COVID-19 and Neurodegeneration.

1. Name potential mechanisms (pathophysiological processes) of Covid-19's impact on the brain described in the text.

2. Describe each process described in the text as semantic triples (subject–predicate–object).

Example:

Pathophysiological Process: Astrocyte_Activation

Triples:

SARS-CoV-2_infection|triggers|astrocyte_activation

If the paragraph does not contain relevant biological content (e.g. acknowledgments, funding, conflicts of interest, publisher notes, metadata, disclaimers, or any non-scientific content), or if no such mechanisms or valid triples are present in the paragraph, return exactly:

Pathophysiological Process: Not_found

Triples:

Not_found|Not_found|Not_found

Use ONLY the information provided in the text! Follow the structure precisely and don't write anything else! Replace spaces in names with _ sign, make sure that words "Pathophysiology Process:" and "Triples:" are presented, don't use bold font and margins. Each triple must contain ONLY THREE elements separated by a | sign, four and more are not allowed!

**S3 Table. GPT-4o prompts used for prompt engineering tasks.**

| **Prompt №** | **Text** |
| --- | --- |
| Prompt_1 | *The URL of an image is given.*  Describe the image (Figure/Graphical abstract) from an article on comorbidity between COVID-19 and Neurodegeneration.  1. Name potential mechanisms (pathophysiological processes) of Covid-19's impact on the brain depicted in the image.  2. Describe each process depicted in the image as semantic triples (subject–predicate–object).  Example:  Pathophysiological Process:  Astrocyte_Activation  Triples:  SARS-CoV-2_infection\|triggers\|astrocyte_activation  Use ONLY the information shown in the image! Follow the structure precisely and don't write anything else! Replace spaces in names with _ sign, make sure that words "Mechanism:" and "Triples:" are presented, don't use bold font and margins. Each triple must contain ONLY THREE elements separated by a \| sign, four and more are not allowed! |
| Prompt_2 | *The URL of an image is given.*  Describe the image (Figure/Graphical abstract) from an article on comorbidity between COVID-19 and Neurodegeneration.  1. Name potential mechanisms (pathophysiological processes) of Covid-19's impact on the brain depicted in the image.  2. Describe each process depicted in the image as semantic triples (subject–predicate–object).  Example:  Pathophysiological Process:  Astrocyte_Activation  Triples:  SARS-CoV-2_infection\|triggers\|astrocyte_activation  Use ONLY the information shown in the image! Follow the structure precisely and don't write anything else! Replace spaces in names with _ sign, make sure that words "Pathophysiological Process:" and "Triples:" are presented, don't use bold font and margins. Each triple must contain ONLY THREE elements separated by a \| sign, four and more are not allowed! The predicate for each triple must be taken only from the Predicate List!  Predicate List: increases, decreases, association, positive_correlation, negative_correlation, regulates, causes_no_change, directly_increases, is_a, directly_decreases, has_variant, biomarker_for, has_members, has_components, orthologous, equivalent_to, has_component, translated_to, prognostic_biomarker_for, has_member, transcribed_to, rate_limiting_step_of, analogous_to.  Structure your answer exactly as it is shown in the example. Don't write anything else! |
| Prompt_3 | *The URL of an image is given.*  Describe the image (Figure/Graphical abstract) from an article on comorbidity between COVID-19 and Neurodegeneration.  1. Name potential mechanisms (pathophysiological processes) of Covid-19's impact on the brain depicted in the image.  2. Describe each process depicted in the image as semantic triples (subject–predicate–object).  Example:  Pathophysiological Process: Astrocyte_Activation  Triples: SARS-CoV-2_infection\|triggers\|astrocyte_activation  Use ONLY the information shown in the image! Follow the structure precisely and don't write anything else! Replace spaces in names with _ sign, make sure that words "Pathophysiology Process:" and "Triples:" are presented, don't use bold font and margins. Each triple must contain ONLY THREE elements separated by a \| sign, four and more are not allowed! The predicates should be chosen according to the provided example, and the objects should represent specific biological elements or conditions.  Predicate examples: increases, decreases, causes, may_cause, leads_to, may_lead_to, affects, may_affect.  Structure your answer exactly as it is shown in the example. Don't write anything else! |

**S4 Table. Full set of semantic triples extracted using GPT-4o prompts and manually curated for evaluation (Can be found in Supplementary_material_S4_Table.xlsx).**
This table contains all semantic triples used in the evaluation of GPT-4o prompt performance. Triples were extracted using three prompt variants (Prompt_1, Prompt_2, Prompt_3) and multiple hyperparameter settings for Prompt_1 (temperature and top_p ranging from 0.0 to 1.0). Additionally, the table includes manually curated gold-standard triples used as reference for similarity scoring (CBM-extracted triples subset). Each row indicates the image number, image URL, Pathophysiological process (if applicable), subject, predicate, and object of the triple.

**S5 Table. Representative semantic comparisons between GPT-predicted and CBM gold-standard triples (image_29)**

|  | **CBM Triple** | **Cosine Similarity** | **Match Status** | **Biological Relevance Summary** |
| --- | --- | --- | --- | --- |
| 1 | inflammatory cytokine increases demyelination | 0.9586 | Match | Canonical downstream effect of cytokine-driven neuroinflammation |
| 2 | inflammatory cytokine increases astrocytosis | 0.9565 | Match | Astrogliosis is a hallmark of CNS inflammation |
| 3 | inflammatory cytokine increases microgliosis | 0.9563 | Match | Microglial activation is central in neuroinflammatory responses |
| 4 | inflammatory cytokine decreases maintenance of blood brain barrier | 0.9408 | Match | BBB disruption is a common cytokine-mediated neuroimmune effect |
| 5 | maternal immune activation increases inflammatory cytokine crossing placenta barrier | 0.9231 | Match | Upstream cause — semantically close and biologically linked via cytokine cascade |
| 6 | increased igg level increases demyelination | 0.9143 | Match | Valid in autoimmune context; linked via neuroinflammatory mechanisms |
| 7 | covid 19 increases maternal immune activation | 0.8600 | Match | Indirectly linked to cytokine release and neuroinflammation (via MIA) |
| 8 | covid 19 increases stress psychological | 0.8218 | No Match | Psychological stress may co-occur but is not mechanistically equivalent |
| 9 | covid 19 increases anxiety | 0.7987 | No Match | Potential downstream outcome, but not semantically or causally equivalent |
| 10 | covid 19 increases depression | 0.7911 | No Match | Correlates weakly with inflammation but not a mechanistic match |

GPT triple: *inflammatory cytokines cause neuroinflammation*
Threshold applied: 0.85 (BioBERT cosine similarity)
Manually inspected GPT-predicted triples were matched against CBM ground truth triples using BioBERT similarity scoring. All shown examples exceeded the threshold of 0.85 and reflect biologically relevant mechanisms linking COVID-19, inflammation, and neurodevelopmental or neurodegenerative processes.

**S6 Table. Performance of GPT-4o prompt variants against CBM gold standard triples.**

| **Prompt** | **Precision** | **Recall** | **F1 Score** | **TP** | **FP** | **FN** |
| --- | --- | --- | --- | --- | --- | --- |
| Prompt_1 | 0.91 | 0.56 | 0.70 | 456 | 43 | 356 |
| Prompt_2 | 0.89 | 0.52 | 0.65 | 400 | 47 | 375 |
| Prompt_3 | 0.96 | 0.48 | 0.64 | 414 | 17 | 448 |

Precision, Recall, and F1-score are computed using BioBERT full-triple matching with a similarity threshold of 0.85. Prompt 1 achieved the best balance of precision and recall, resulting in the highest F1-score. Prompt 3 had the highest precision but lower recall, while Prompt 2 showed the weakest overall performance.

**S7 Table. Impact of temperature and top_p on Prompt_1 extraction performance.**

| **Setting** | **Precision** | **Recall** | **F1 Score** | **TP** | **FP** | **FN** |
| --- | --- | --- | --- | --- | --- | --- |
| Temp = 1.0, Top_p = 1.0 | 0.93 | 0.49 | 0.65 | 389 | 29 | 398 |
| Temp = 0.75, Top_p = 0.75 | 0.91 | 0.54 | 0.67 | 405 | 41 | 349 |
| Temp = 0.5, Top_p = 0.5 | 0.92 | 0.55 | 0.69 | 446 | 38 | 363 |
| Temp = 0.25, Top_p = 0.25 | 0.90 | 0.54 | 0.68 | 440 | 47 | 369 |
| **Temp = 0.0, Top_p = 0.0** | **0.91** | **0.56** | **0.70** | **456** | **43** | **356** |

Precision, recall, and F1 scores for GPT-4o triples generated with Prompt_1 under different decoding parameter settings, compared to the CBM gold standard using BioBERT full-triple matching (threshold = 0.85).

### **Supplementary figures**

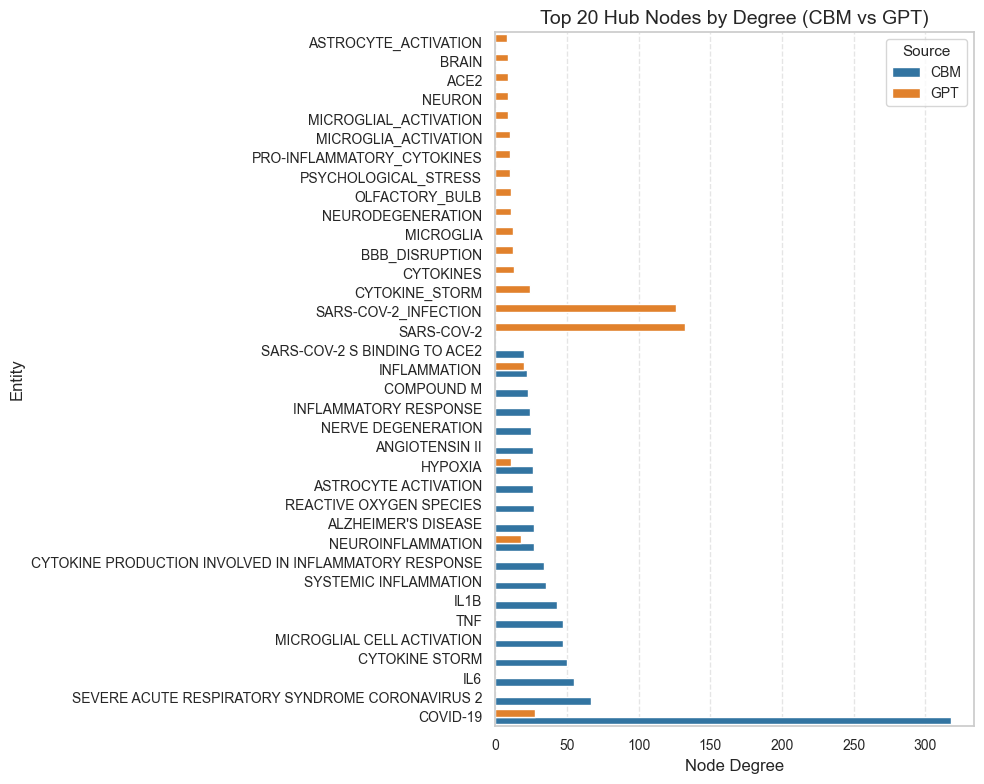

**Fig. S1. Top 20 hub nodes by degree in CBM-curated and GPT-extracted knowledge graphs.**

Comparison of top 20 hub nodes (most connected entities) between CBM‑curated and GPT‑extracted semantic graphs. Horizontal bars represent node degree (number of incident edges). CBM nodes are shown in green; GPT nodes are shown in orange. The hub nodes highlight frequently occurring biomedical entities in each knowledge extraction modality.

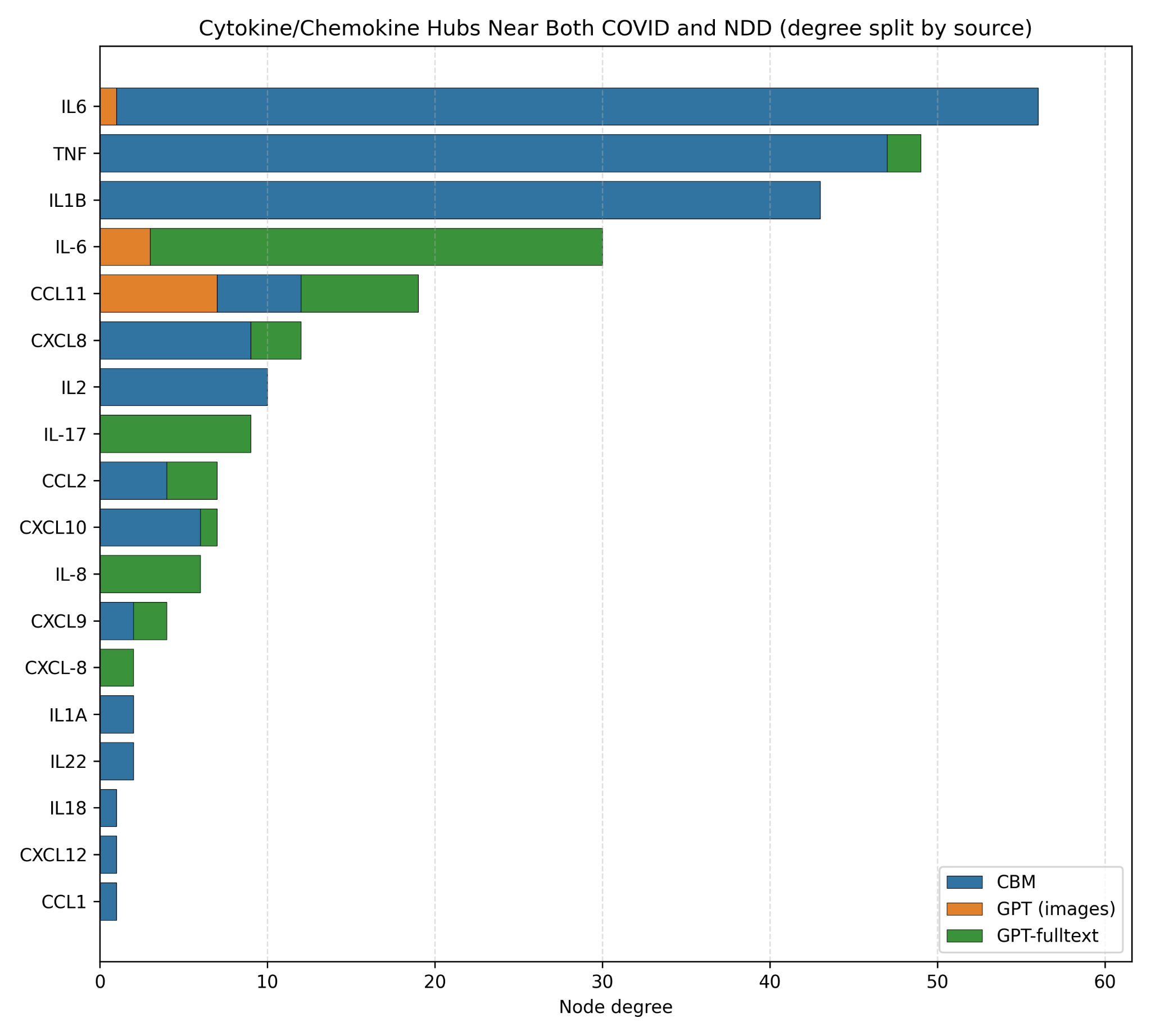

**Fig. S2. Cytokine/chemokine hubs near both COVID-19 and NDD.**

A subset restricted to cytokines/chemokines. CBM emphasizes IL-6, TNF, and IL-1β, while GPT adds CXCL10, CXCL12, CCL2, and IL-17, expanding the inflammatory repertoire.

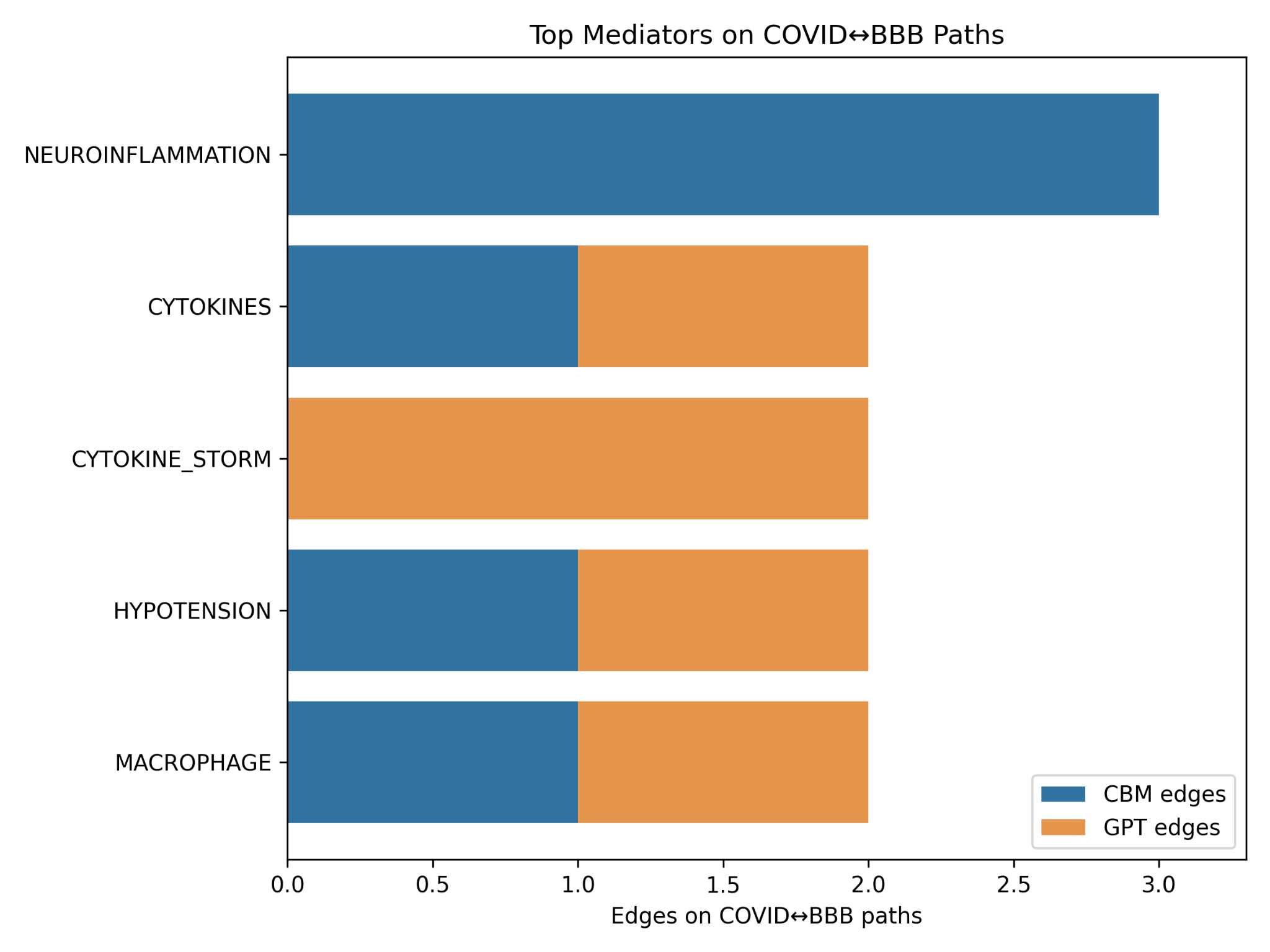

**Fig. S3. Top mediators on COVID↔BBB shortest paths.**

Stacked bar plot showing mediators linking COVID and BBB disruption. CBM strongly emphasizes neuroinflammation; GPT adds systemic factors (cytokine storm, hypotension, macrophages).

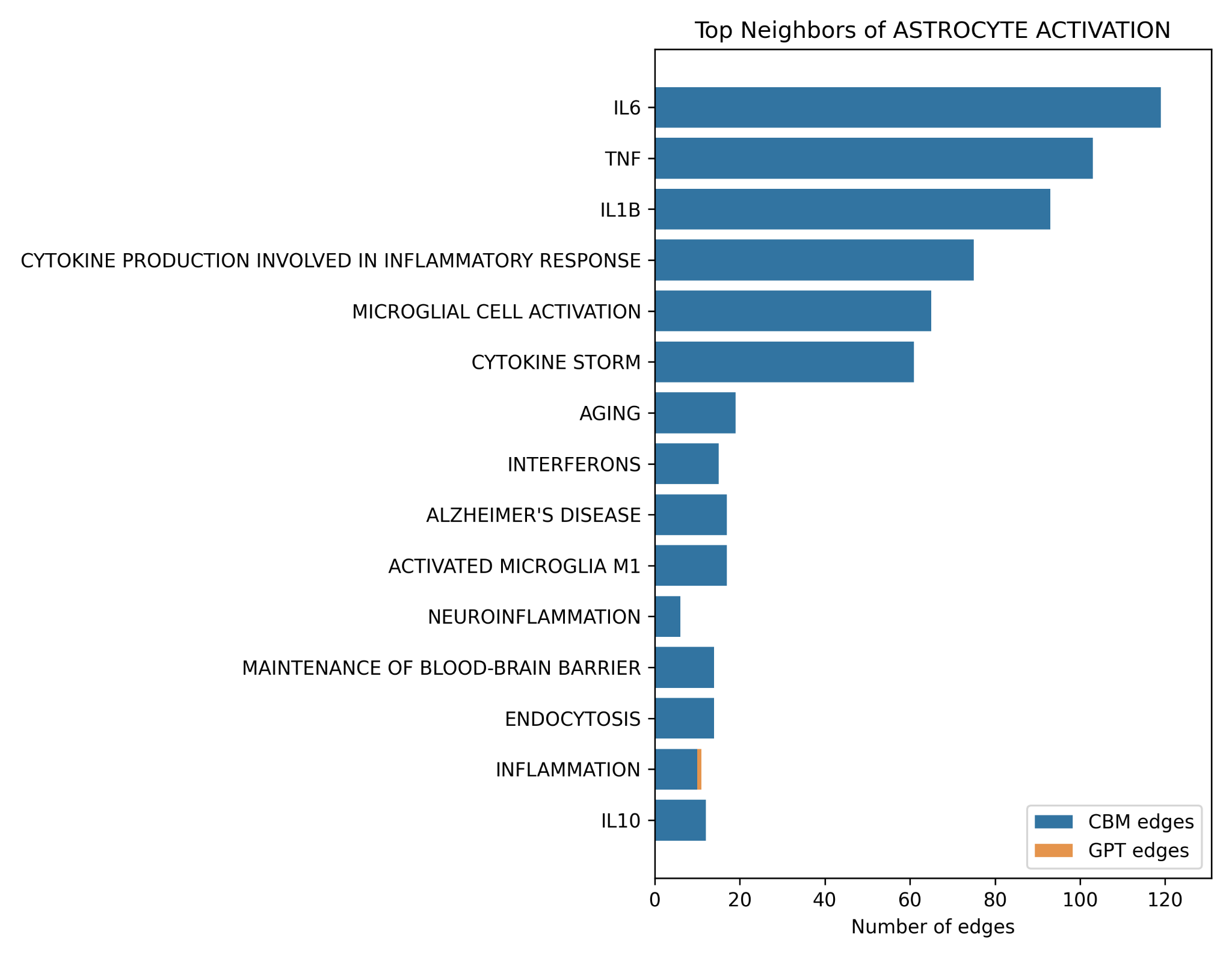

**Fig. S4. Top neighbors of astrocyte activation.**

Top 15 neighboring nodes of astrocyte activation. CBM emphasizes IL-6, TNF, and IL-1β, while GPT contributes oxidative stress, autoantibodies, and chemokines.
